# GEI-17-Mediated SUMOylation of GDI-1 Regulates Apoptotic Cell Clearance

**DOI:** 10.1101/2025.01.22.634223

**Authors:** Lei Yuan, Peiyao Li, Aiying Ma, Chao Li, Fuqiao Liu, Yunmin Xie, Qian Zheng, Hui Wang, Hui Xiao

## Abstract

Apoptotic cell clearance represents the final stage of apoptosis, and involves phagosome formation, maturation, and digestion. Here, we found that SUMO modifications participate in apoptotic cell clearance by influencing phagosomal degradation activity in *C. elegans*. The SUMO E3 ligase, GEI-17, SUMOylates GDP dissociation inhibitor (GDI-1) at K270 to trigger phagosomal degradation. Upon SUMOylation, GDI-1 facilitates release of GDP-bound RAB-1, which is subsequently converted to the GTP-bound form in a manner dependent on GDI displacement factor, PRAF-3. RAB-1 conversion enables binding and translocation of RAB-7 from the endoplasmic reticulum to Golgi to promote phagosome maturation and degradation activity. In the absence of GDI-1 SUMOylation, GDI-1 binds to GDP-RAB-1, sequestering it in the cytoplasm, resulting in impaired phagosomal degradation. Furthermore, we found SUMOylation of GDI1 at a conserved site plays a crucial role in efferocytosis regulation in mammals. This study defines a previously unrecognized mechanism by which SUMOylation drives apoptotic cell clearance.

## Introduction

Apoptosis plays an important role in the development of multicellular organisms and maintenance of homeostasis in tissue (Boada-Romero, Martinez et al., 2020, Doran, Yurdagul et al., 2020, Schilperoort, Ngai et al., 2023). As the final step in apoptosis, cell corpse clearance is a highly dynamic process involving mutual recognition between apoptotic cells and phagocytic receptors, cytoskeletal rearrangement in phagocytes, phagosome formation in the phagocytes extending pseudopods to engulf apoptotic cells, and ultimately phagosomal degradation within phagocytes (Kawano & Nagata, 2018, Nagata, 2018, Yang & Wang, 2021). Defects in apoptotic cell clearance have been implicated in the development of various severe chronic inflammatory or catastrophic autoimmune disorders (e.g., systemic lupus erythematosus) as well as neurodegenerative diseases (e.g., Alzheimer’s disease) (Poon & Ravichandran, 2024, Romero-Molina, Garretti et al., 2022).

During development of *C. elegans*, 131 somatic cells and approximately 50% of germ cells undergo apoptosis, and apoptotic cells are subsequently phagocytosed and degraded by neighboring cells, such as body wall muscle cells and gonadal sheath cells (Wang & Yang, 2016, Xie, Chen et al., 2024). Apoptotic cell clearance is evolutionarily highly conserved, and apoptotic cells are degraded by phagocytosis by two parallel and partially redundant pathways. One is composed of CED-1/MEGF10, CED-6/GULP and CED-7/ABC transporter membrane proteins (Chen, Xiao et al., 2010, Mapes, Chen et al., 2012, Zhou, Hartwieg et al., 2001), and the other is composed of CED-2/CrkII, CED-5/Dock180, CED-10/Rac and CED-12/Elmo proteins (Reddien & Horvitz, 2000, Wang, Wu et al., 2003). Under the synergistic action of these two signaling pathways, the pseudopods of phagocytosis are extended to engulf apoptotic cells to form phagosomes (Fazeli, Levin-Konigsberg et al., 2023, Lee, Hamann et al., 2019). Subsequently, phagosomes mature in the presence of a series of Rab GTP hydrolases (RAB-5, RAB-2, RAB-14, and RAB-7), and the mature phagosomes fuse with lysosomes to form phagolysosomes. Finally, apoptotic cellular components within the phagolysosome are degraded by lysosomal hydrolases (Guo, Hu et al., 2010, Li, Zou et al., 2009, Poteryaev, Fares et al., 2007).

Ubiquitin and SUMOylation are very important post-translational modifications of proteins, are implicated in various biological processes, including DNA damage repair, cell proliferation and apoptosis, etc (Li, Jing et al., 2021, Qin, Li et al., 2021, Zheng & Shabek, 2017). Our previous research has demonstrated that in *Drosophila melanogaster*, the E3 ubiquitin ligase Pallbearer is involved in phagocytosis by ubiquitinating RpS6, resulting in the activation of the RAC2 GTPase, which subsequently leads to the upregulation of F-actin remodeling (Xiao, Wang et al., 2015). Our recent studies have revealed that the E3 ubiquitin ligase TRIM-21 and the E2 conjugating enzyme UBC-21 function synergistically to catalyze K48-mediated polyubiquitination modification of the phagocytosis receptor CED-1 in *C. elegans*, targeting proteasomal degradation and promoting apoptotic cell clearance (Yuan, Li et al., 2022a). These studies demonstrated that ubiquitination modifications are very important for apoptotic cell clearance. Whereas ubiquitylation is primarily associated with the protein turnover, SUMOylation primarily facilitates or inhibits protein interactions and alters protein conformation or localization, especially with proteins that contain SUMO-interacting motifs (SIMs), thereby enabling transient binding to target proteins (Vertegaal, 2022, Zhao, 2018). SUMOylation is a reversible process that is conserved in *C. elegans*. Analogous to ubiquitination, SUMOylation occurs through the action of an E1-activating enzyme (the Sae1/Sae2 heterodimer in humans and AOS-1/UBA-2 in worms), an E2-conjugating enzyme (Ubc9 in humans and UBC-9 in worms), and SUMO-specific E3 ligases (PIAS and MMS21 in humans and GEI-17 and MMS-21 in worms) (Pelisch, Tammsalu et al., 2017, Princz, Pelisch et al., 2020). However, some SUMOylation reactions do not involve E3 ligases. The deconjugation of SUMO from its targets is regulated by SUMO-specific isopeptidases (SENP1, 2, 3, 5, 6, and 7 in humans and ULP-1, ULP-2, ULP-4, and ULP-5 in worms) (Drabikowski, Ferralli et al., 2018, Tsur, Bening Abu-Shach et al., 2015). Mammals contain three different SUMO proteins, whereas, *C. elegans* expresses only a single SUMO orthologue, SMO-1, which renders the worm an expedient model organism to dissect this post-translational protein modification system. SUMOylation has been predominantly investigated in the development in *C. elegans*, and has been implicated in embryonic, vulval, and muscle development, as well as in DNA damage response (Fergin, Boesch et al., 2022, Kim, Ding et al., 2021b, Princz et al., 2020). Disturbances in SUMOylation are closely related to the pathogenesis of a variety of human diseases such as neurodegenerative diseases, diabetes mellitus, innate immune diseases and tumors. Compared to the study of ubiquitination modification in apoptotic cell clearance, the study of SUMO modification mostly focuses on the process of apoptosis. However, it remains unclear whether SUMO modifications are involved in the clearance of apoptotic cells.

To determine whether and how SUMO modifications contribute to apoptotic cell clearance, we induced SUMO pathway knockdown in *C. elegans* and found that cell corpse degradation, lysosomal recruitment of LAAT-1, NUC-1, and CPL-1, and phagosomal acidification were all impaired. Proteomic analysis revealed that GDP dissociation inhibitor 1 (GDI-1) was the substrate for SUMO modification required for apoptotic cell clearance, specifically at residue K270. In addition, through numerous genetic and chemical manipulation approaches, we found that GDI-1 SUMOylation by the E3 ligase, GEI-17, modulated cell corpse removal via regulation of RAB-1 activity through a mechanism dependent on competing for binding with the GDI displacement factor, PRAF-3, in *C. elegans*. Specifically, SUMOylated GDI-1 promotes RAB-1 binding and transport of RAB-7 from the endoplasmic reticulum to Golgi apparatus to positively regulate apoptotic cell clearance. In addition, we found that the human homolog of GEI-17, PIAS1, functioned similarly in regulating phagocytosis in mammalian cells, and that it could reciprocally complement GEI-17 activity towards *C. elegans* proteins *in vitro*. Moreover, we found that SUMOylation of GDI1 also occurs in mammals, where it contributes to the regulation of efferocytosis. These results suggesting that this SUMOylation-dependent apoptotic cell clearance pathway may be conserved between nematodes and humans. Our study thus defines a previously undocumented regulatory mechanism for cell corpse clearance via regulation of GDI-1 SUMOylation, suggesting possible therapeutic targets for further exploration to alleviate apoptosis-associated diseases.

## Results

### The SUMO system participates in cell corpse clearance in *C. elegans*

To determine whether and how SUMOylation may participate in regulating apoptosis in *C. elegans*, we induced silencing of *smo-1* by RNA interference (RNAi). Examination of cell corpse numbers showed that worms with *smo-1* knockdown had higher numbers of cell corpses both in the germline, in an age-dependent manner, and in embryos at six developmental stages from comma to 4F compared to that in the non-targeted RNAi controls (Fig. 1A-C). As *ced-3* and *ced-4* are key factors in apoptotic cell death in *C. elegans,* loss-of-function (lf) mutations in *ced-3* or *ced-4,* caused an almost complete loss of apoptotic cell death (Yuan & Horvitz, 1990). Subsequent examination of cell corpses showed that apoptotic cell death was almost completely lost in the absence of either gene. Furthermore, *ced-3(n717)* or *ced-4(n1126)* mutants treated with *smo-1* RNAi also had significantly fewer apoptotic germ cells (Fig. 1D), suggesting that the germ cell corpses we observed in *smo-1* knockdown animals resulted from physiological cell death. As the *C. elegans* SUMO modification system also includes the E1 activator, AOS-1/UBA-2, the E2 conjugating enzyme, UBC-9, and the E3 ligase homologs, GEI-17/MMS-21, we also induced knockdown of individual SUMOylation pathway components, and found that the number of apoptotic germline cells significantly increased compared to control (Fig. 1E).

**Figure 1.**
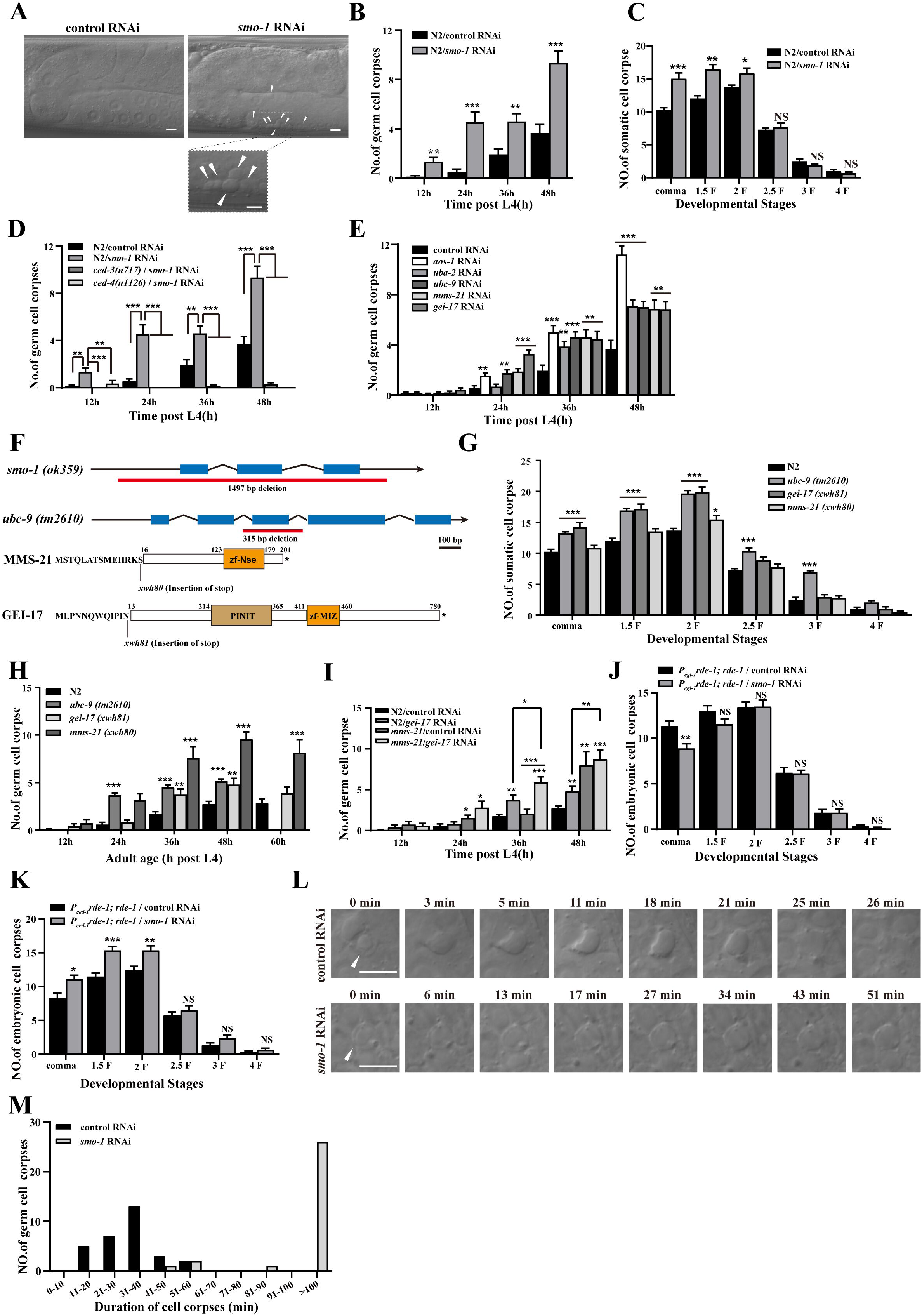
The SUMO system regulates apoptotic cell clearance in *C. elegans*. (A) DIC (differential interference contrast) images of N2 treated with the control or *smo-1* RNAi germline. Arrows indicate button-like cell corpses. The boxed region is magnified (2.5×) in the inset. Bars, 5 µm. (B and C) Quantification (mean ± SEM) of button-like cell corpses at different adult stages (h post L4) (B) and different embryonic stages (C) in N2/control and *smo-1* RNAi worms. Fifteen adult worms and embryos were scored at each stage for each strain. (D-E and G-K) The development of adult or embryo stages corpses were quantified (mean ± SEM) in the indicated strains. Fifteen adult worms or embryos were scored at each stage for each strain. (F) Schematic illustration of the *smo-1 (ok359)*, *ubc-9 (tm2610)*, *mms-21 (xwh80)*, and *gei-17 (xwh81)* homozygous mutations generated by CRISPR-Cas9 editing of the endogenous *mms-21* and *gei-17* loci. Amino acids near the mutated sites are indicated. (L) Time-lapse recording of cell corpses in N2/control and *smo-1* RNAi germlines. Arrows indicate cell corpses. Bars, 5 µm. (M) Four-dimensional microscopy analysis of the persistence of 30 germ cell corpses duration were performed in N2/control and *smo-1* RNAi. The unpaired *t* test was performed in this figure. *P < 0.05, **P < 0.01, ***P < 0.001, NS, no significance. All error bars indicate means and SEM.

To further investigate how the SUMO system affects programmed cell death in *C. elegans*, we analyzed *smo-1(ok359)* and *ubc-9(tm2610)* deletion mutants, which represent strong loss-of-function mutations (Fig. 1F). In addition, we used CRISPR-Cas9 to generate homozygous mutant alleles of the SUMO E3 ligases, *gei-17(xwh81)* and *mms-21(xwh80),* carrying nonsynonymous point mutations leading to premature N-terminal stop codons (Fig. 1F). The *smo-1(ok359)* mutation resulted in an embryonic apoptosis phenotype that precluded examination of cell corpses due to complete early embryonic lethality. However, we observed significantly increased numbers of cell corpses at multiple embryonic stages and in *ubc-9(tm2610)*, *gei-17(xwh81)*, and *mms-21(xwh80)* adult germline cells relative to that in WT controls (Fig. 1G,H). Notably, *gei-17(xwh81)* mutants had higher embryonic cell corpse numbers than *mms-21(xwh80)* mutants, whereas more germline cell corpses were detected in the *mms-21(xwh80)* mutant than *gei-17(xwh81)*. Although unable to generate homozygous *gei-17(xwh81); mms-21(xwh80)* double mutants, we obtained *gei-17(xwh81); mms-21(xwh80)* double heterozygotes, potentially due to redundant functions of these genes required for viability and normal development in *C. elegans.* Alternatively, RNAi knock-down of *gei-17* resulted in significantly enhanced numbers of germ cell corpses in the *mms-21* mutant, suggesting that the two E3 ligases might belong to independent pathways (Fig. 1I). Taken together, these results demonstrated that SUMO modifications contributed to regulating apoptosis in *C. elegans*.

To investigate whether the increase in apoptotic cells was due to activation of apoptosis in a higher proportion of cells or defects in apoptotic cell clearance, we treated *xwhIs49(P_ced-1_rde-1; rde-1)* transgenic worms (with phagocyte-specific RNAi suppression) and *xwhIs82(P_egl-1_rde-1; rde-1)* transgenic worms (with apoptotic cell-specific RNAi suppression) with *smo-1* RNAi, *gei-17* RNAi, and *mms-21* RNAi. We found that cell corpse counts significantly increased in *P_ced-1_rde-1; rde-1,* but not *P_egl-1_rde-1; rde-1,* worms compared to their respective RNAi vector controls, in resulted in increased cell corpse counts in (Figs. 1K and EV1A,B), suggesting that silencing *smo-1*, *gei-17*, and *mms-21* affected cell corpse removal rather than induction of apoptosis. We next conducted four-dimensional microscopy to measure the duration of embryonic or germ cell corpse persistence. In wild-type animals, the germ cell corpses were seldom persisted for >60 min. By contrast, *smo-1* knockdown led to significantly longer persistence in gonads for the large majority of germ cell corpses (>100 min) (Fig. 1L,M). In addition, germ cell corpses persisted significantly longer in *gei-17(xwh81)* and *mms-21(xwh80)* mutants than in WT worms (*Figure S1C*). Additionally, cell corpse removal required significantly more time in *gei-17(xwh81) and mms-21(xwh80)* embryos compared to wild-type embryos (Fig. EV1D,F). However, the number of cell deaths in *gei-17(xwh81) and mms-21(xwh80)* embryos was indistinguishable from that in the wild type (Fig. EV1E,G). These findings indicated that SUMO modifications participated in the clearance of apoptotic cells rather than induction of apoptosis.

As phagocytosis of apoptotic cells is controlled by two partially redundant signaling pathways, CED-1/CED-6/CED-7/NRF-5/TTR-52/DYN-1 and CED-2/CED-5/CED-10/CED-12/PSR-1, in *C. elegans*, we next examined whether SUMO modifications functioned conjointly with engulfment genes to regulate cell corpse clearance. To this end, we treated the *ced-1(e1735)* and *ced-6(n1813)* engulfment mutants with *smo-1* RNAi and found that *smo-1* silencing led to significantly enhanced defects in germ cell corpse engulfment at different adult stages in both mutants (Fig. EV1H). Intriguingly, we found that *smo-1* RNAi treatment also resulted in greater impairment of germ cell corpse engulfment in *ced-5(n1812)* and *ced-12(n3261)* mutants, the alternate regulatory pathway for cell corpse engulfment (Fig. EV1H). Additionally, *smo-1* RNAi treatment also led to increased germ cell corpse abundance in the *ced-1(e1735); ced-2(n1994)* double mutant (Fig. EV1I). These findings suggested that SUMO modifications functioned independently of either pathway for cell corpse removal.

### SUMOylation promotes apoptotic cell clearance through phagosome maturation

Since phagosome assembly and subsequent clearance of apoptotic cells involves several steps in *C. elegans,* we investigated the possible role of SUMOylation in phagosomal recruitment of factors required for phagosome maturation. Fluorescence microscopy with a CED-1::GFP phagocytic receptor fusion reporter in germline cells of *smo-1* knockdown worms indicated that cell corpses were surrounded by CED-1::GFP, similar to vector controls (Fig. 2A,G), which suggested that recognition and initiation of engulfment were unaffected by *smo-1* suppression. To determine whether SUMOylation affects phagosomal recruitment, we examined GFP-labeled markers for different stages of phagosome maturation, including GFP::RAB-5, a small GTPase specific to early endosomes and phagosomes; GFP::RAB-7, a small GTPase required for late endosome–phagosome fusion (Kinchen, Doukoumetzidis et al., 2008); LAAT-1::GFP, a lysosomal membrane protein (Liu, Du et al., 2012); NUC-1::mCHERRY, a lysosomal DNase (Lyon, Evans et al., 2000); and CPL-1::mChOint, a lysosomal cathepsin protease (Xu, Liu et al., 2014). We found that the percentage of phagosomes surrounded by GFP::RAB-5 and GFP::RAB-7 in germ cells did not significantly differ between *smo-1* RNAi-treated worms and vector controls (Fig. 2B,C,G), whereas significantly fewer phagosomes were surrounded with LAAT-1::GFP under *smo-1* knockdown (Fig. 2D,G). Moreover, NUC-1::mCHERRY and CPL-1::mChOint signals were also significantly decreased on phagosomes in the germ line of *smo-1* knockdown animals compared to that in control animals (Fig. 2E,F,G). These data suggested that SUMO modifications regulate recruitment to phagosome in the later stages of maturation.

**Figure 2.**
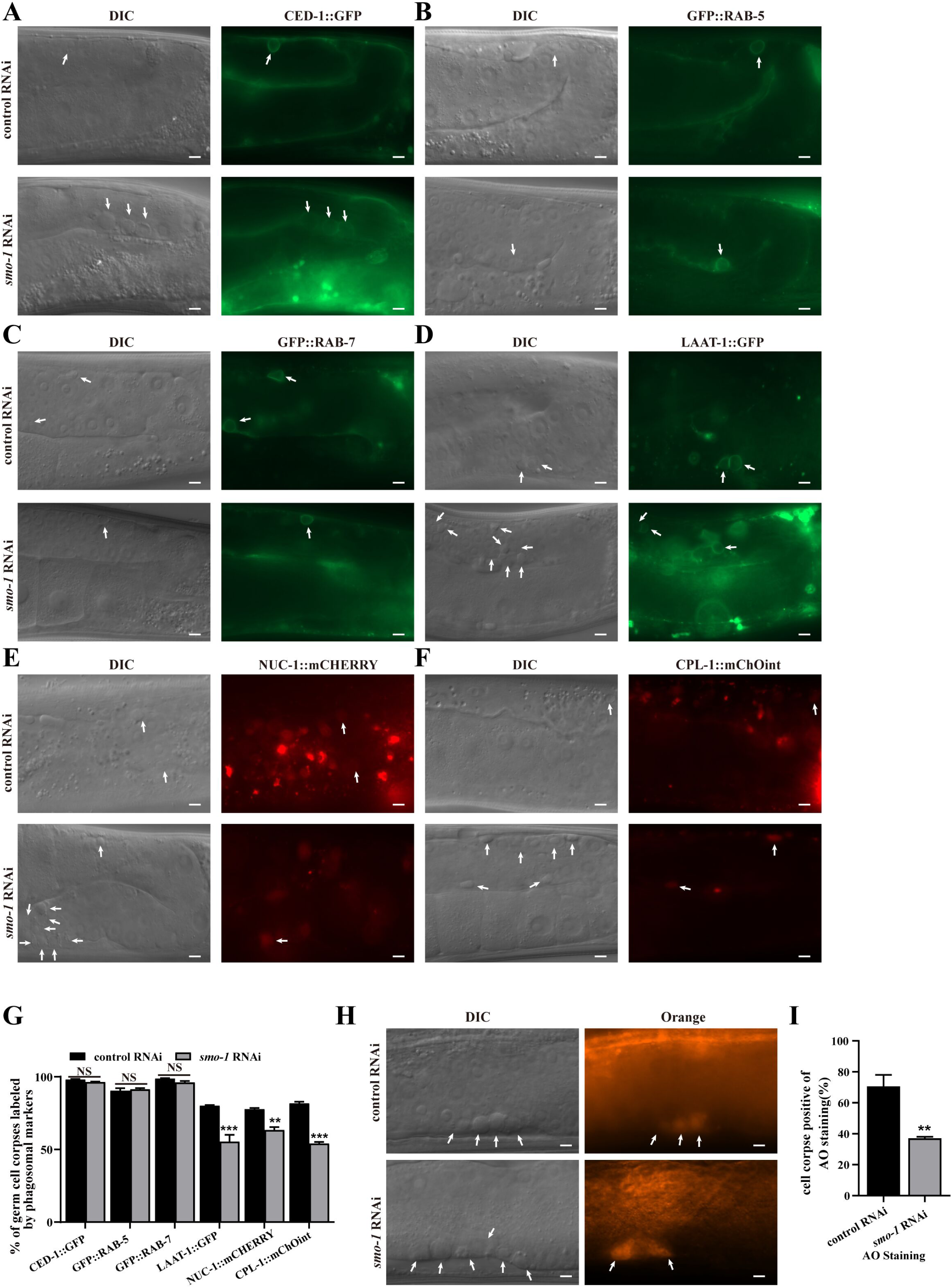
SUMOylation promotes apoptotic cell clearance through phagosome maturation. (A-F) Representative images of cell corpse labeling by CED-1::GFP (A), GFP::RAB-5 (B), GFP::RAB-7 (C), LAAT-1::GFP (D), NUC-1::mCHERRY (E), and CPL-1::mChOint (F). Cell corpses are indicated by arrows. Bars, 5 μm. (G) Quantification (mean ± SEM) of the labeling of cell corpses by different phagosomal markers shown in A–F. ≥100 cell corpses were analyzed for each marker; and data were derived from three replicates. Comparisons were performed between the N2/control and *smo-1* RNAi worms for each marker. (H) AO staining of germ cell corpses from the indicated animals. Arrows indicate cell corpses in the N2/control and *smo-1* RNAi worms. Bars, 5 μm. (I) Quantification (mean ± SEM) of AO-positive germ cell corpses in N2/control and *smo-1* RNAi worms. The unpaired *t* test was performed in this figure. **P < 0.01, ***P < 0.001, NS, no significance. All error bars indicate the mean ± SEM.

To test whether the persistence of cell corpses in *smo-1* RNAi-treated worms was due to failed acidification of phagosomes containing cell corpses, a late stage of corpse clearance, acridine orange (AO) staining was used to identify compromised cell corpses. This analysis indicated that *smo-1* (RNAi) worms had obviously decreased AO staining compared with control RNAi worms (Fig. 2H,I), supporting the likelihood that impaired SUMO modification affected the acidification of phagosomes following cell corpse engulfment.

To further examine the impact of SUMOylation on post-engulfment phagosomal degradation, we next quantified endosomal/lysosomal degradation activity in *smo-1* knockdown worms using *arIs36(P_heat-shock_ssGFP)* transgenic worms that expressed ssGFP under the control of a heat-shock-inducible promoter. Monitoring of ssGFP uptake and degradation by coelomocytes over 48 hours showed that ssGFP began to accumulate in the body cavity (pseudocoelom) of control RNAi worms within one hour post-heat shock (hphs) treatment, and was subsequently transported to coelomocytes (Fig. EV2A). The ssGFP signal then progressively decreased in the body cavity over 12 hphs, coinciding with increased GFP signal in coelomocytes. At 24 hphs, ssGFP was no longer detectable in the body cavity or coelomocytes (Fig. EV2A), indicating that ssGFP was taken in and degraded by coelomocytes. In *smo-1* knockdown animals, ssGFP was transported to coelomocytes by 1 hphs, similar to that in controls, suggesting that SUMOylation does not affect ssGFP uptake into coelomocytes. However, ssGFP was still detectable in the body cavity as well as coelomocytes of *smo-1* RNAi-treated animals at 48 hphs (Fig. EV2A), indicating that endosomal/lysosomal protein degradation was markedly impaired in coelomocytes of the *smo-1* knockdown group. These results suggested that suppressing SUMOylation resulted in obviously attenuated endosomal/lysosomal degradation activity.

To directly observe the process of cell corpse degradation, we next induced *smo-1* knockdown in transgenic *ujIs113*(*P_pie-1_H2B::mCherry; P_nhr-2_HIS-24::mCherry*) worms expressing germline-specific markers of chromatin H2B::mCHERRY and HIS-24::mCHERRY. We observed that chromatin was condensed in germ cell corpses and subsequently disappeared by around 30 minutes (31.00±2.83 min, n=5). On the contrary, we observed that chromatin was condensed in germ cell corpses in *smo-1* RNAi worms, but then diffused throughout the corpse, rather than disappearing, in later stage phagosomes, with mCHERRY signal persisting for >70 min (80.00±7.94 min, n=5) (Fig. EV2B,C). These results suggested chromatin degradation was significantly delayed in the cell corpse-containing phagosomes of *smo-1* knockdown worms. Collectively, these findings demonstrated that cell corpse accumulation associated with *smo-1* knockdown was due to defects in cell corpse degradation in engulfing cells.

### GDI-1 is the SUMOylation substrates potentially involved in apoptotic cell clearance

To identify the SUMOylation substrates potentially involved in apoptotic cell clearance in *C. elegans*, we generated an *xwh83*(*ha::smo-1*) worm strain by CRISPR-Cas9 gene editing that expressed *smo-1* with an N-terminal HA tag as its sole copy of the SUMO gene (Fig. EV3A). We then immunoprecipitated (IP) SUMO-conjugated proteins with an anti-HA antibody and identified components of the isolated protein mixture by LC-MS/MS analysis (Fig. 3A). Given the widespread expression of SUMO in *C. elegans,* and our use of a mixed worm population, we anticipated identifying targets across all stages of development and tissues. In total, this analysis uncovered 188 SUMOylated proteins in *C. elegans* under normal developmental conditions (Table 1).

**Figure 3.**
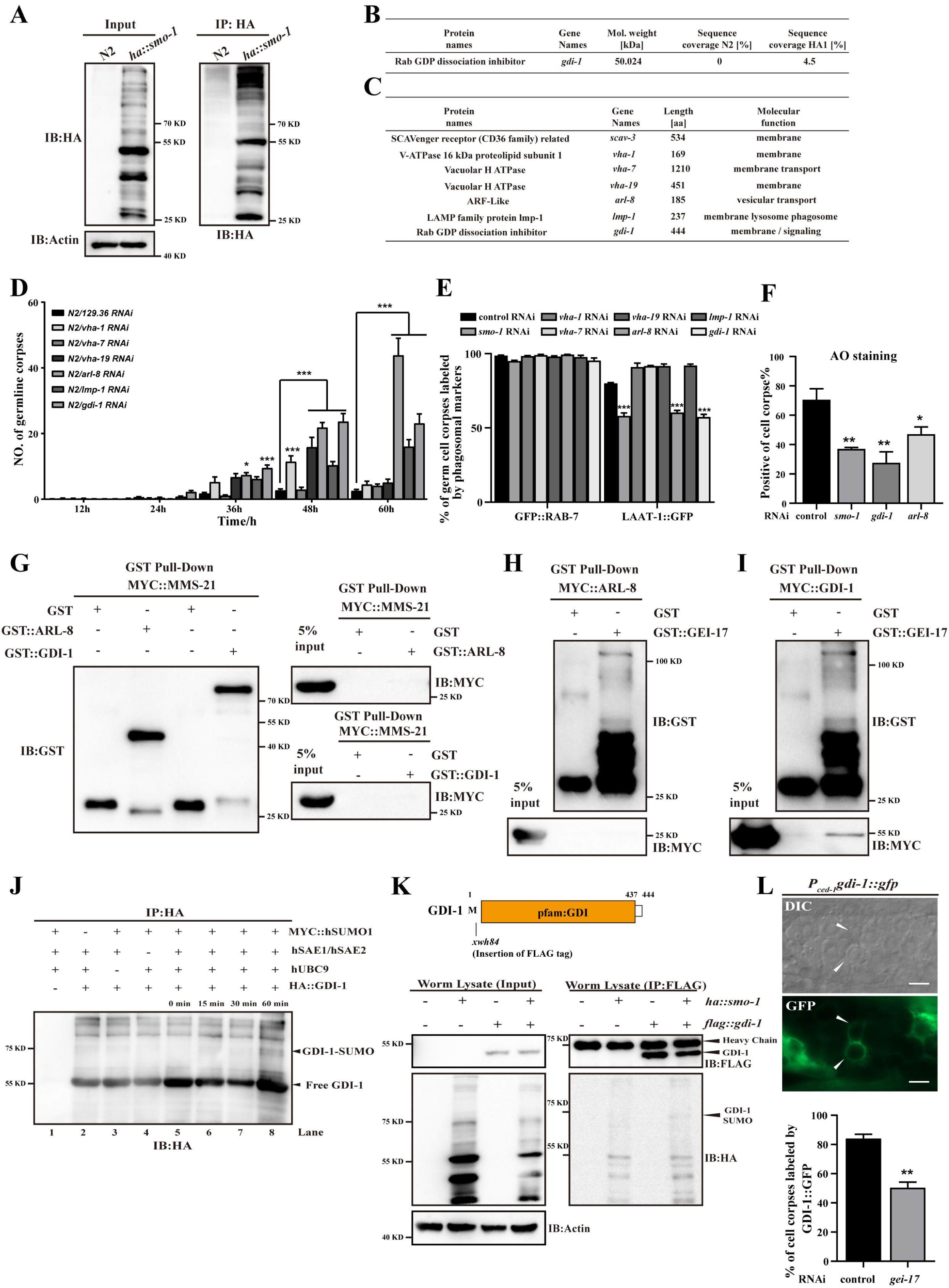
GDI-1 is the SUMOylation substrates potentially involved in apoptotic cell clearance. (A) HA IP was performed on *ha::smo-1* worms, followed by the detection of proteins that interact with SMO-1 using LC-MS/MS. IB, immunoblot. (B) The potential SMO-1 interacting proteins were identified by protein mass. (C) A list of potential substrates for SUMOylation associated with lysosomes and vesicles, with data derived from the protein mass spectrometry results of SUMO-GFP. (D) Quantification (mean ± SEM) of germ cell corpses from the indicated strains. Cell corpses of each animal were scored in fifteen animals at every time point (h post L4) as indicated. (E) Quantification (mean ± SEM) of cell corpses positive for phagosomal markers in indicated strains. A total of ≥100 cell corpses were analyzed for each marker; and data were derived from three replicates. (F) Quantification (mean ± SEM) of AO-positive germ cell corpses in the N2/control, *smo-1*, *gdi-1*, and *arl-8* RNAi worms. (G-I) The interactions between MMS-21-ARL-8/MMS-21-GDI-1 (G), GEI-17-ARL-8 (H), and GEI-17-GDI-1 (I) were examined by GST pull-down assays. (J) Recombinant HA::GDI-1 is modified by hSUMO1 *in vitro*. Recombinant HA::GDI-1 was incubated for the indicated times (minutes) in modification reactions, including E1 (hSAE1 and hSAE2), E2 (hUBC9), hSUMO1, and ATP (lanes 5–8). HA IP was performed, followed by detection of SUMOylation with anti-HA antibodies. SUMOylation of GDI-1 was observed (lane 8 was compared with lanes 1–7). (K) SUMOylation of GDI-1 was examined in N2, *ha::smo-1*, *flag::gdi-1*, and *ha::smo-1; flag::gdi-1*. FLAG IP was performed, followed by detection of SUMOylation with anti-HA antibodies. (L) Representative images (top) and quantification (bottom) of GDI-1::GFP localization on cell corpses in control and *gei-17* RNAi germlines. Arrows indicate cell corpses. Bars, 5 µM. The unpaired *t* test was performed in this figure. *P < 0.05, **P < 0.01, ***P < 0.001. All error bars indicate the mean ± SEM.

**Table 1.**
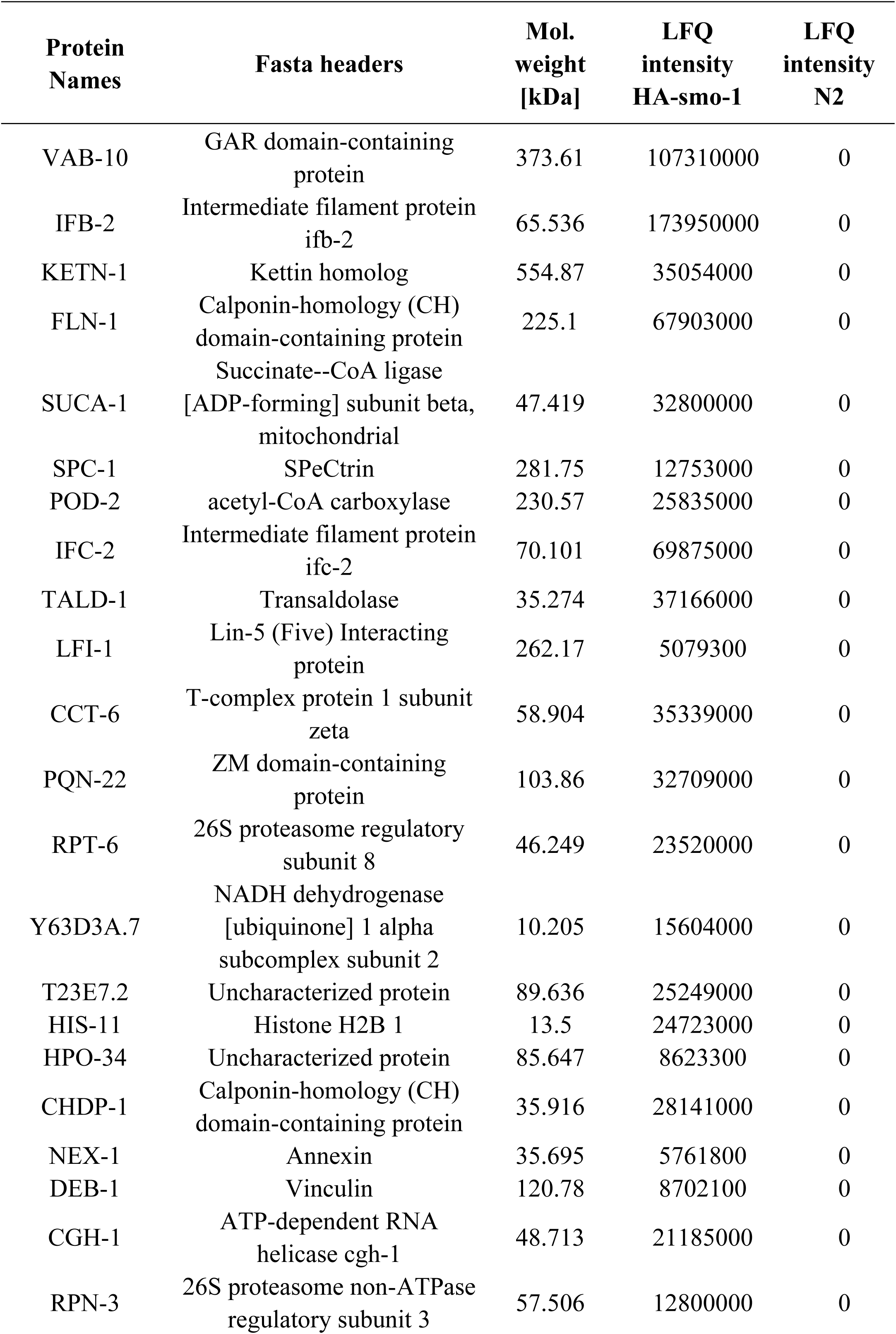
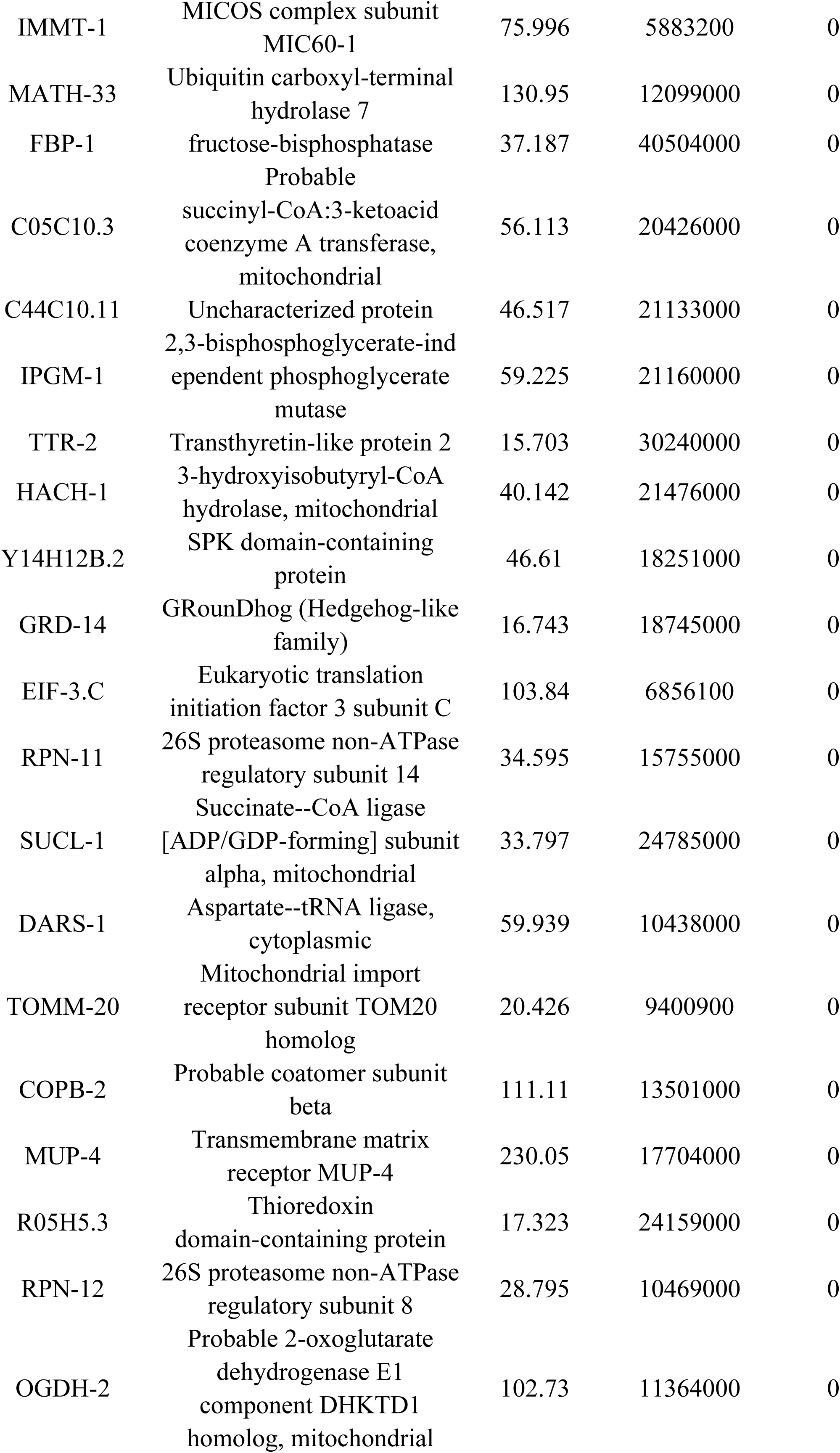
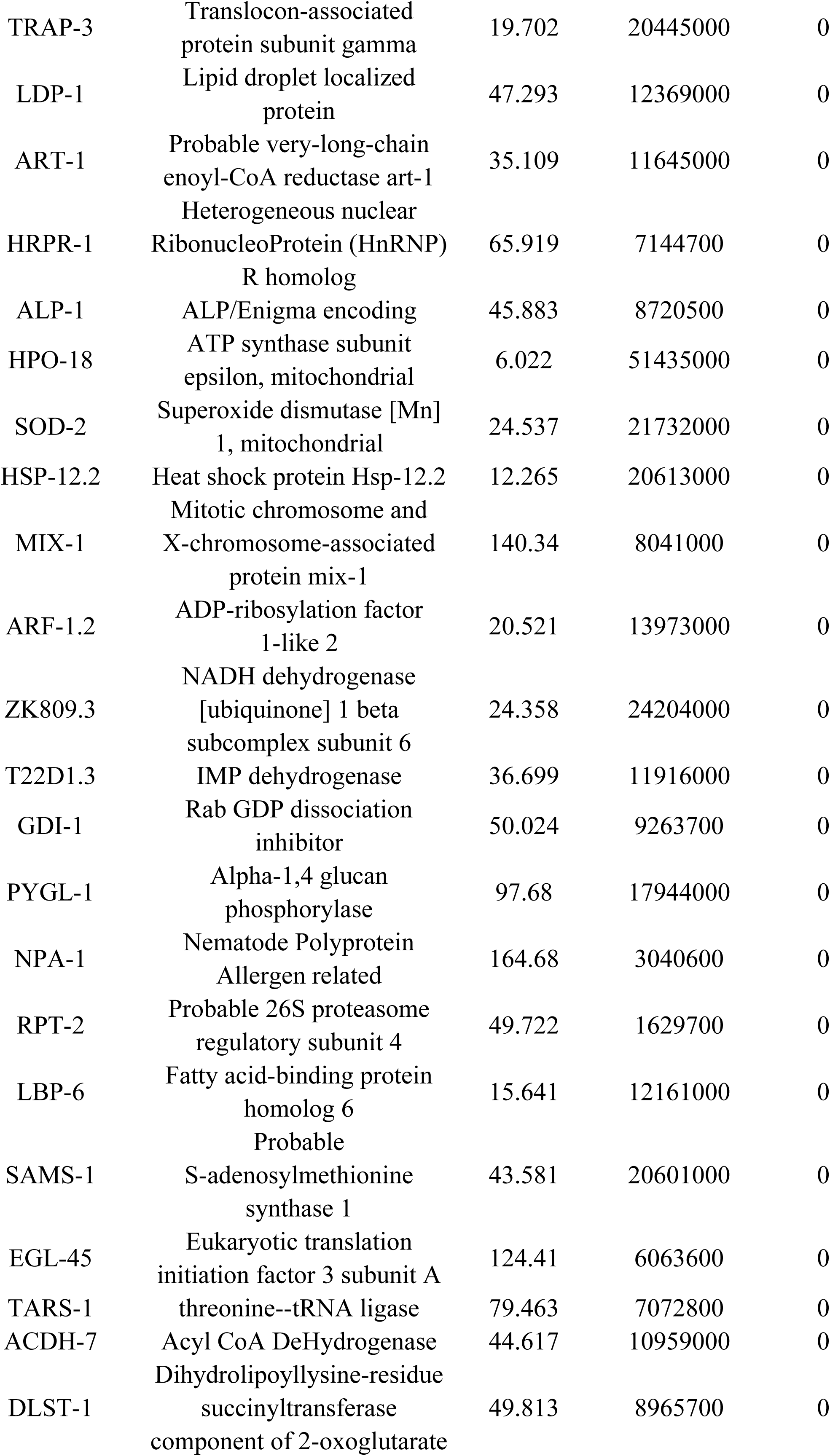
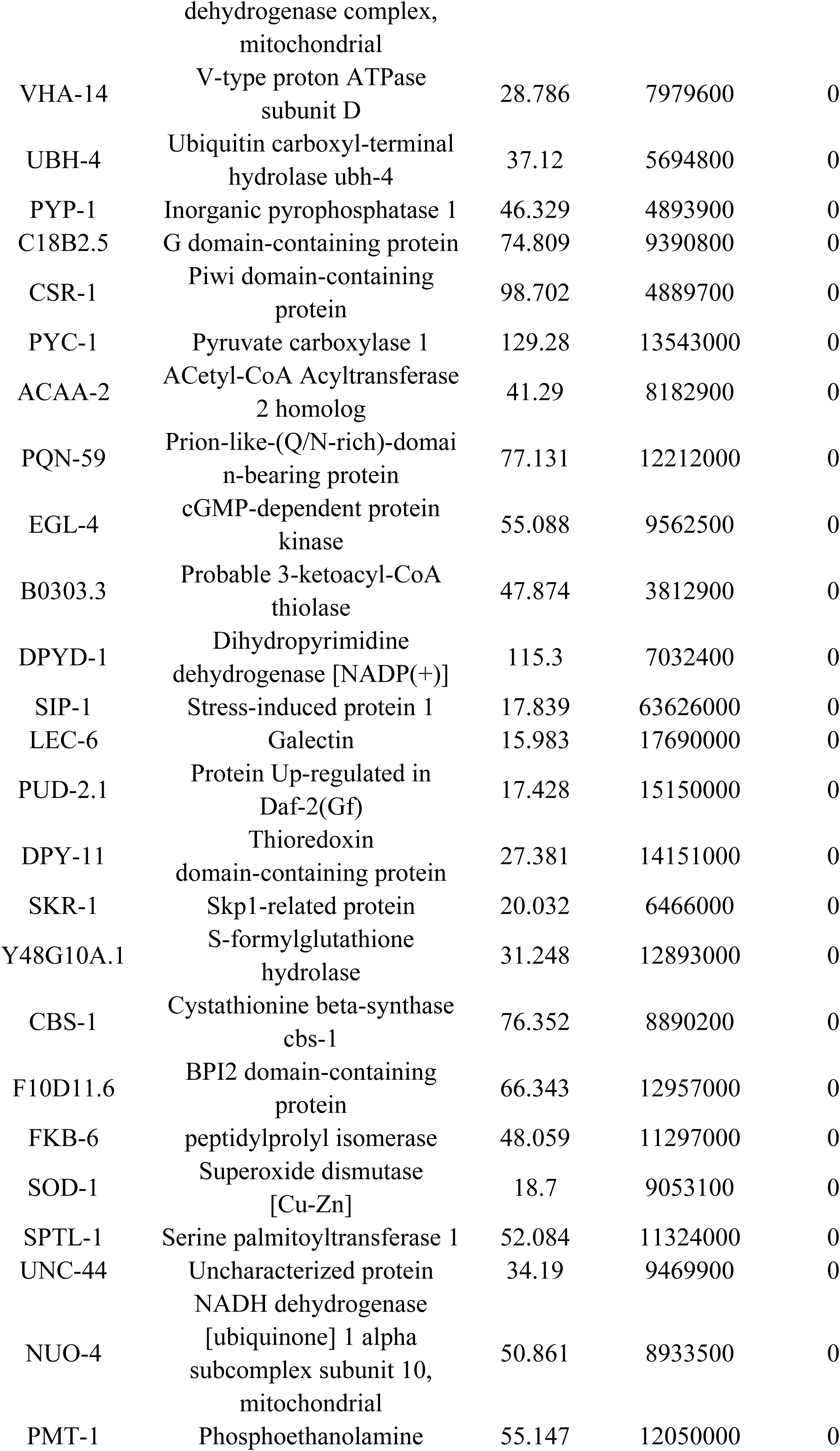
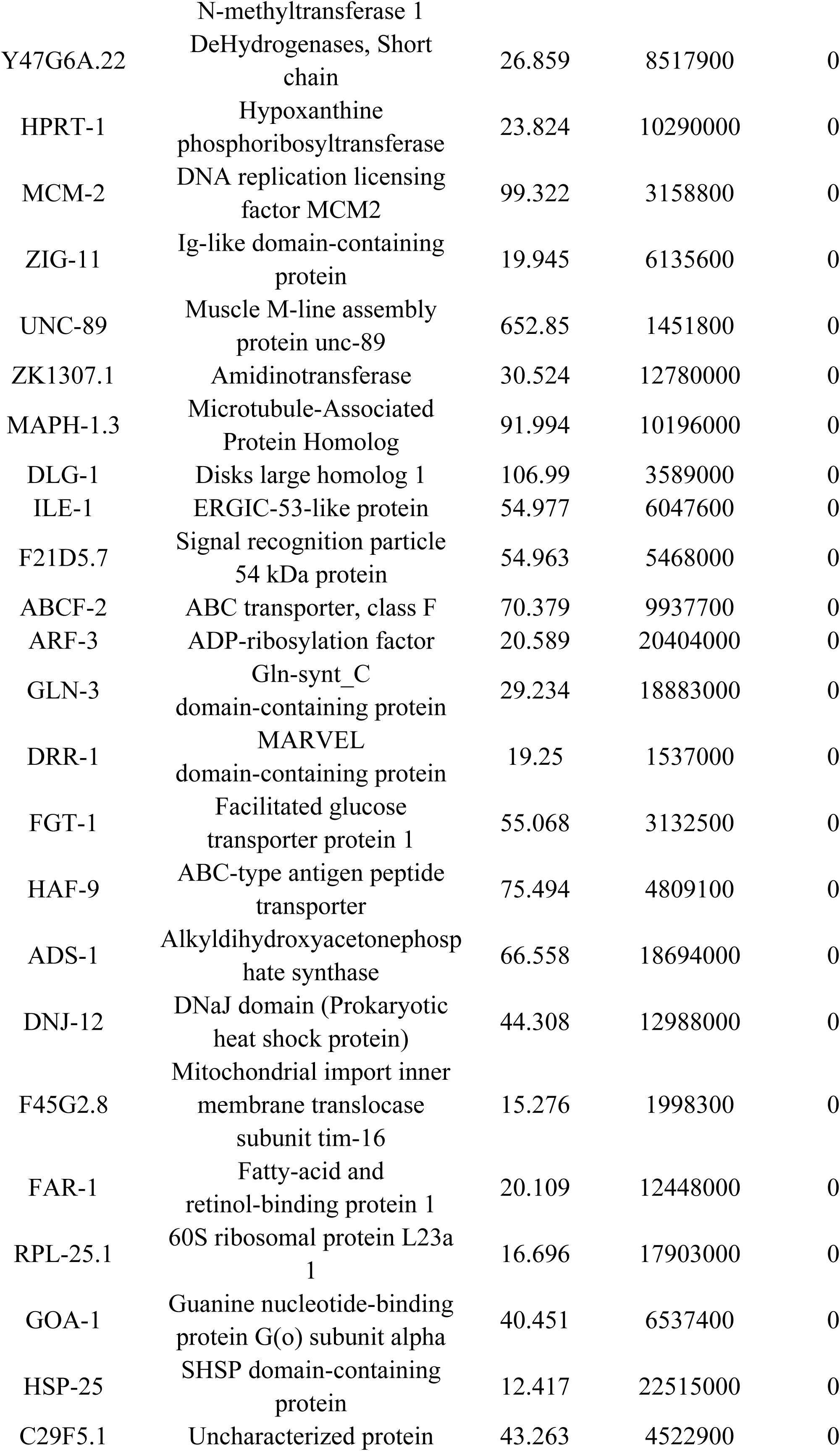
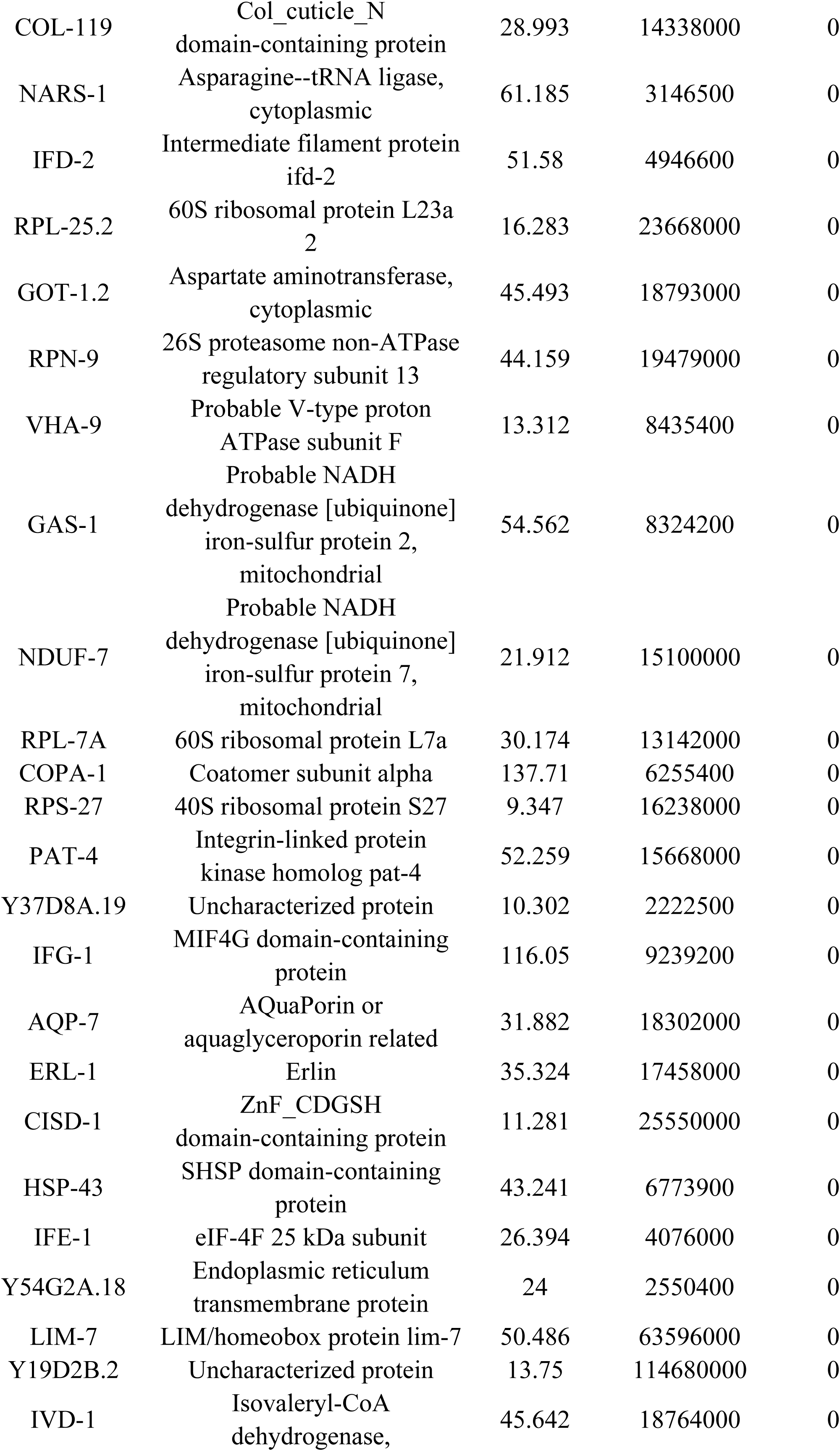
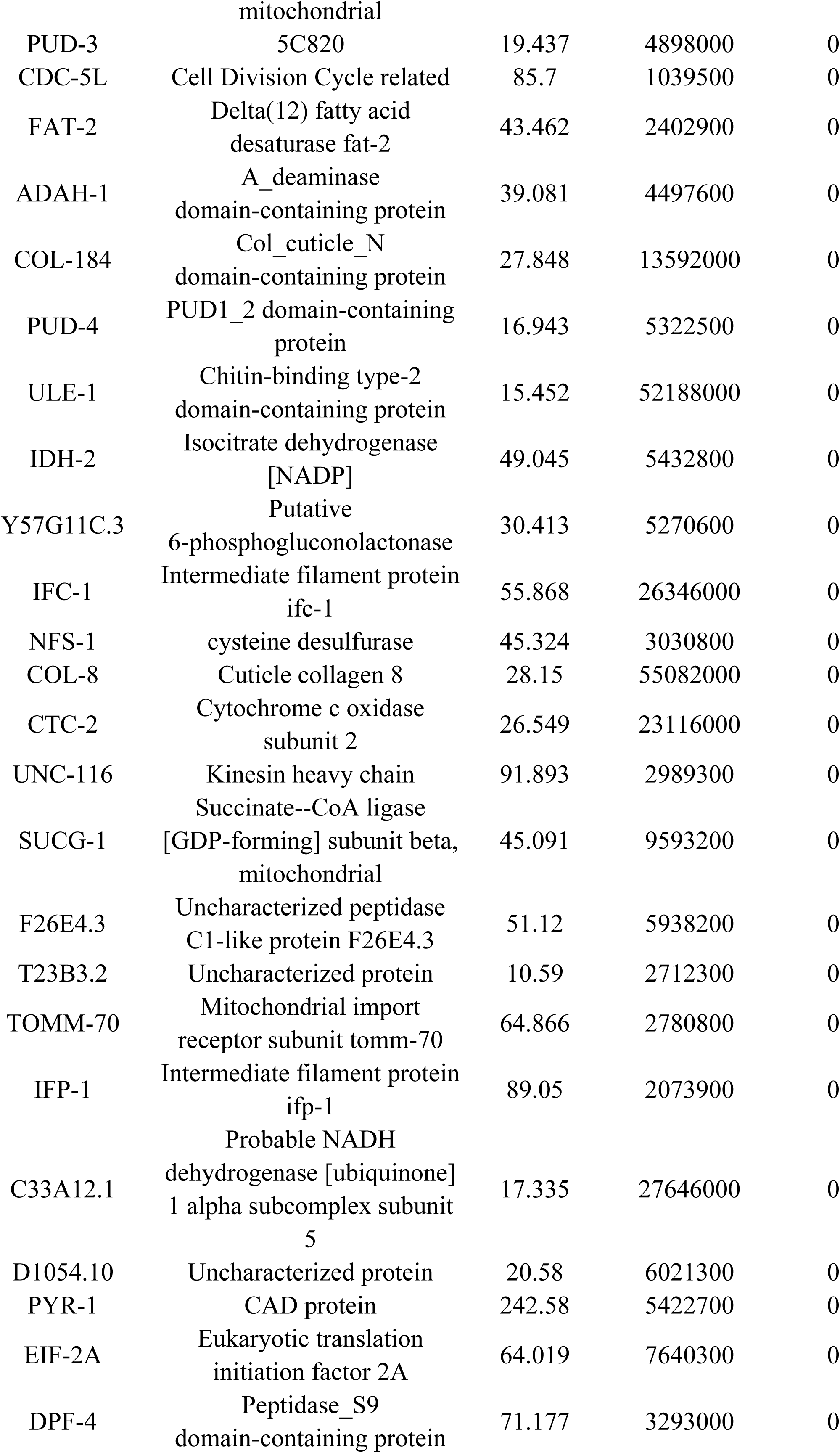
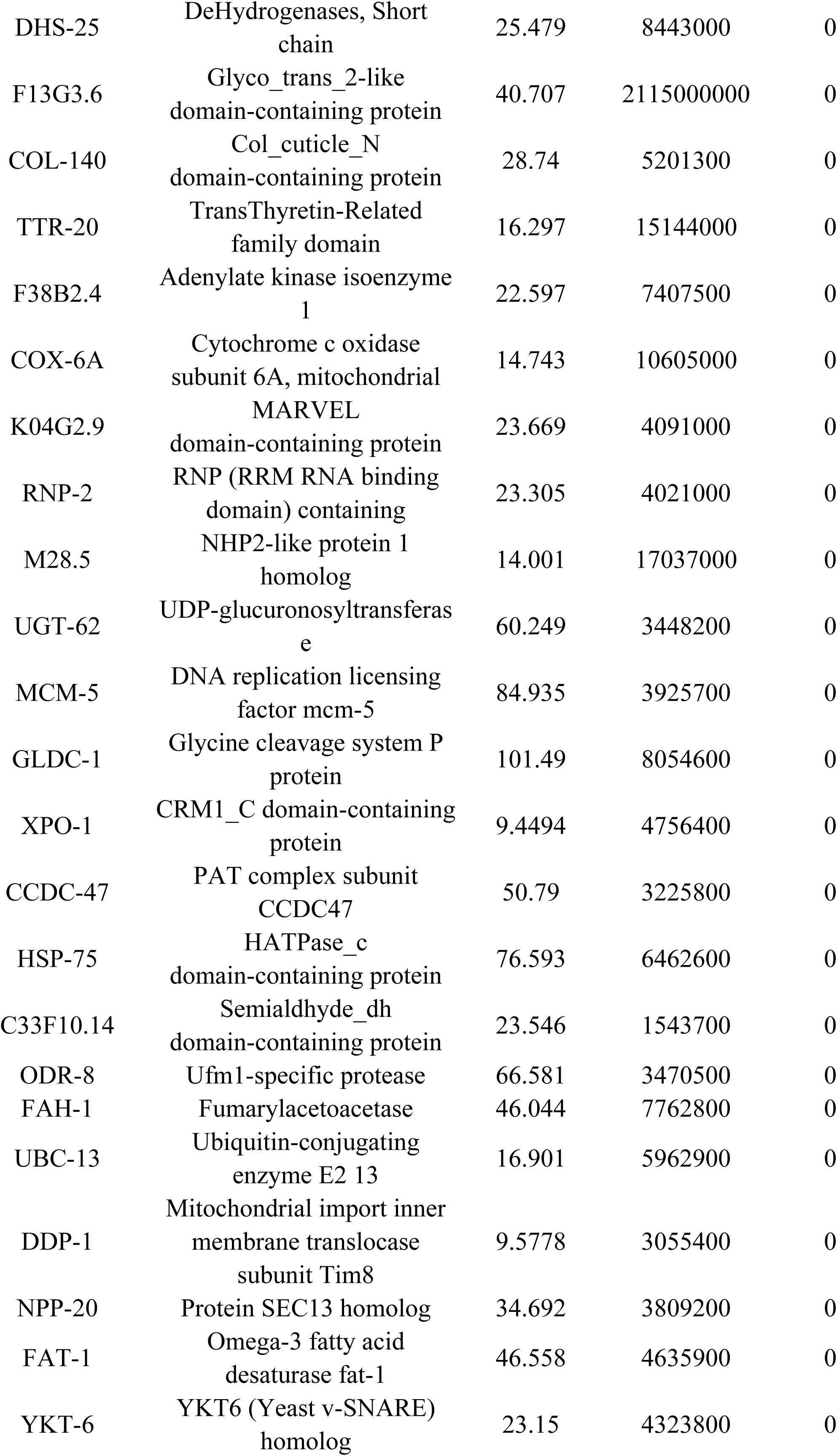
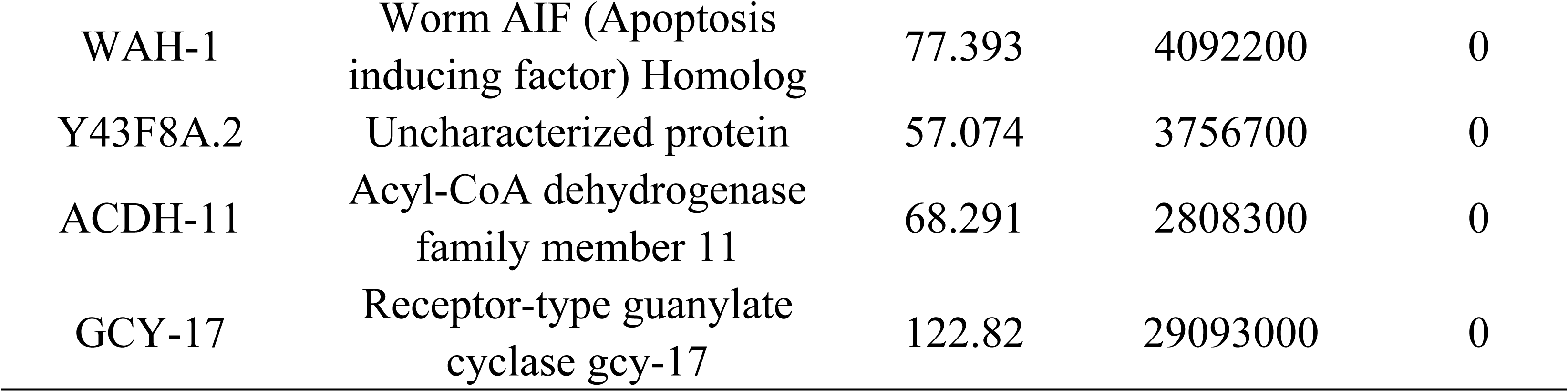
The potential SUMOylated proteins identified by LC-MS/MS in *C. elegans*.

We then examined Gene Ontology (GO) functional annotations of the SUMO-modified proteins (Fig. EV3B), and selected the RAB GDP dissociation inhibitor, *gdi-1*, due to its reported involvement in vesicular transport (Fig. 3B), and based on our understanding that SUMO likely promotes apoptotic cell clearance through phagosome maturation in our above experiments. In addition to *gdi-1*, we examined six other candidates identified in similar proteomics studies in *C. elegans* and mammalian cells (Becker, Barysch et al., 2013, Drabikowski et al., 2018) (Fig. 3C), including LMP-1, a membrane protein found in cytoplasmic vesicles and lysosomal membranes; ARL-8, a small lysosomal GTPase that regulates axonal dendritic transport, phagosome-to-lysosome translocation, and synaptic vesicle paracrine transport; and three VHA family members that regulate lysosomal luminal acidification through proton-transporting ATPase activity. Among these candidates, inactivation of *arl-8* or *gdi-1* by RNAi in WT worms produced an apoptotic phenotype similar to *smo-1* knockdown (Fig. 3D-F).

To further elucidate the SUMO substrate involved in apoptotic cell clearance, we first examined ARL-8 and GDI-1 interactions with SMO-1 by Yeast two-hybrid assays (Y2H), and found that both ARL-8 and GDI-1 likely exhibited interactions with SMO-1. As SMO-1 can undergo direct covalent or non-covalent binding with substrates, or may interact with substrates indirectly through covalent or non-covalent binding with an intermediary interaction partner, we generated SMO-1 truncation variants with C-terminal deletions that either exposed the conserved bis-glycolide motif (GG’: active form) or deleted the bis-glycine motif (ΔGG: inactive form) (Gao, Li et al., 2019). Subsequent Y2H assays showed that WT and SMO-1 GG’, but not the SMO-1 ΔGG variant, were capable of interacting with ARL-8 or GDI-1 (Fig. EV3C). These results suggested that SMO-1 exhibited covalent, rather than non-covalent, direct or indirect interactions to modify ARL-8 or GDI-1. Co-IP and pull-down assays including two SUMO E3 ligases and ARL-8 or GDI-1 suggested that GDI-1 could exclusively interact with GEI-17, but not MMS-21, whereas, ARL-8 did not interact with either E3 ligase (Figs. 3G-I and EV3D). These results suggested that GDI-1 might serve as substrate of GEI-17 [which has been shown to interact with SMO-1]. We therefore focused on GEI-17 function in apoptotic cell clearance.

To identify the regions of GDI-1 required for interaction with GEI-17, we generated a series of GDI-1 truncation constructs based on secondary structures identified in its amino acid sequence. Co-IP and GST pull-down assays showed that only the region between residues 1-150 of GDI-1 was required for binding with GEI-17 (Fig. EV3E-G). As SUMO-1/2/3 serve as precursor molecules in mammals, while SAE1/2 function as E1 activating enzymes, and UBC9 acts as the E2 binding enzyme, we next tested whether GDI-1 was a SUMO substrate through *in vitro* SUMOylation reactions using the human homologs of these proteins expressed and purified in *E. coli*. We found that recombinant hUBC9, hSAE1/hSAE2, and hSUMO1 could together SUMOylate GDI-1 (Figs. 3J and EV3H). Notably, the SUMOylation of GDI-1 was completely abolished by excluding either hSAE1/hSAE2 or hUBC9 (Fig. 3J), indicating that GDI-1 was a bona fide SUMOylation target *in vitro*.

To determine whether GDI-1 was SUMOylated in *C. elegans in vivo*, we inserted a flag tag into the endogenous *gdi-1* loci using CRISPR-Cas9, which resulted in the *xwh84(flag::gdi-1) C. elegans* strain (Fig. 3K). We then crossed the *xwh83*(*ha::smo-1*) and *xwh84(flag::gdi-1)* strains to obtain progeny carrying both tagged proteins and performed anti-FLAG IP to test whether GDI-1 was directly modified with SMO-1. Western blot detection of HA in IP lysates of the *flag::gdi-1; ha::smo-1* recombinant progeny revealed a high-molecular mass species (Fig. 3K) that likely represented SUMOylated GDI-1 in these worms.

To next investigate whether and how SUMO modification of GDI-1 was involved apoptotic cell clearance in *C. elegans,* we generated a transgenic *xwhIs85(P_gdi-1_gdi-1::gfp)* strain expressing GDI-1::GFP driven by the native *gdi-1* promoter. Interestingly, fluorescence microscopy showed that GDI-1::GFP was mainly present in pharyngeal and body wall muscle cells (Fig. EV3I). Since apoptotic cells are engulfed by neighboring cells in *C. elegans*, including hypodermal cells, intestinal cells, muscle cells, and gonadal sheath cells, together with our finding that GDI-1 was mainly localized in pharyngeal and body wall cells, we hypothesized that GDI-1 could play a role in phagocytosis of neighboring cell corpses. To test this possibility, we constructed a transgenic *xwhIs86(P_ced-1_gdi1::gfp)* strain expressing GDI-1::GFP driven by the native *ced-1* promoter. Subsequent fluorescence microscopy analysis showed that germ cell corpses were surrounded by GDI-1::GFP in the *xwhIs86* worms. Moreover, we observed that significantly lower levels of GDI-1::GFP labeled germline corpses were recruited to phagosomes in *gei-17* RNAi-treated worms compared to that in vector control animals (Fig. 3L). These data suggested that SUMO modification of GDI-1 by GEI-17 indeed participated in apoptotic cell clearance and likely played an important role in the cell corpse degradation process.

### SUMOylation of GDI-1 at K270 is required for apoptotic cell degradation

In order to identify the specific lysine residue on GDI-1 targeted for SUMOylation by the E3 SUMO ligase, we crossed the *xwh83*(*ha::smo-1*) and *xwhIs86*(*P_ced-1_gdi-1::gfp)* strains to obtain progeny co-expressing both tagged proteins for anti-HA co-IP assays. Western blot detection of HA in IP lysates of the recombinant progeny revealed a smear of high-molecular mass protein species (Fig. 4A). Subsequent LC-MS/MS of the purified lysates (Table 2) identified K113, K175, and K270 as predicted SUMO-acceptor sites on GDI-1 through SUMOplot analysis of the MS protein profile (Fig. 4B). Site-directed mutagenesis converting the candidate lysine sites to arginines resulted in single, double, and triple GDI-1 SUMOylation site mutants, including K113R, K113R/K175R, K113R/K175R/K270R. Y2H analysis of potential interaction between these mutants and SMO-1 showed that only the GDI-1(K270R) lost the ability to interact with SMO-1 (Fig. 4C).

**Figure 4.**
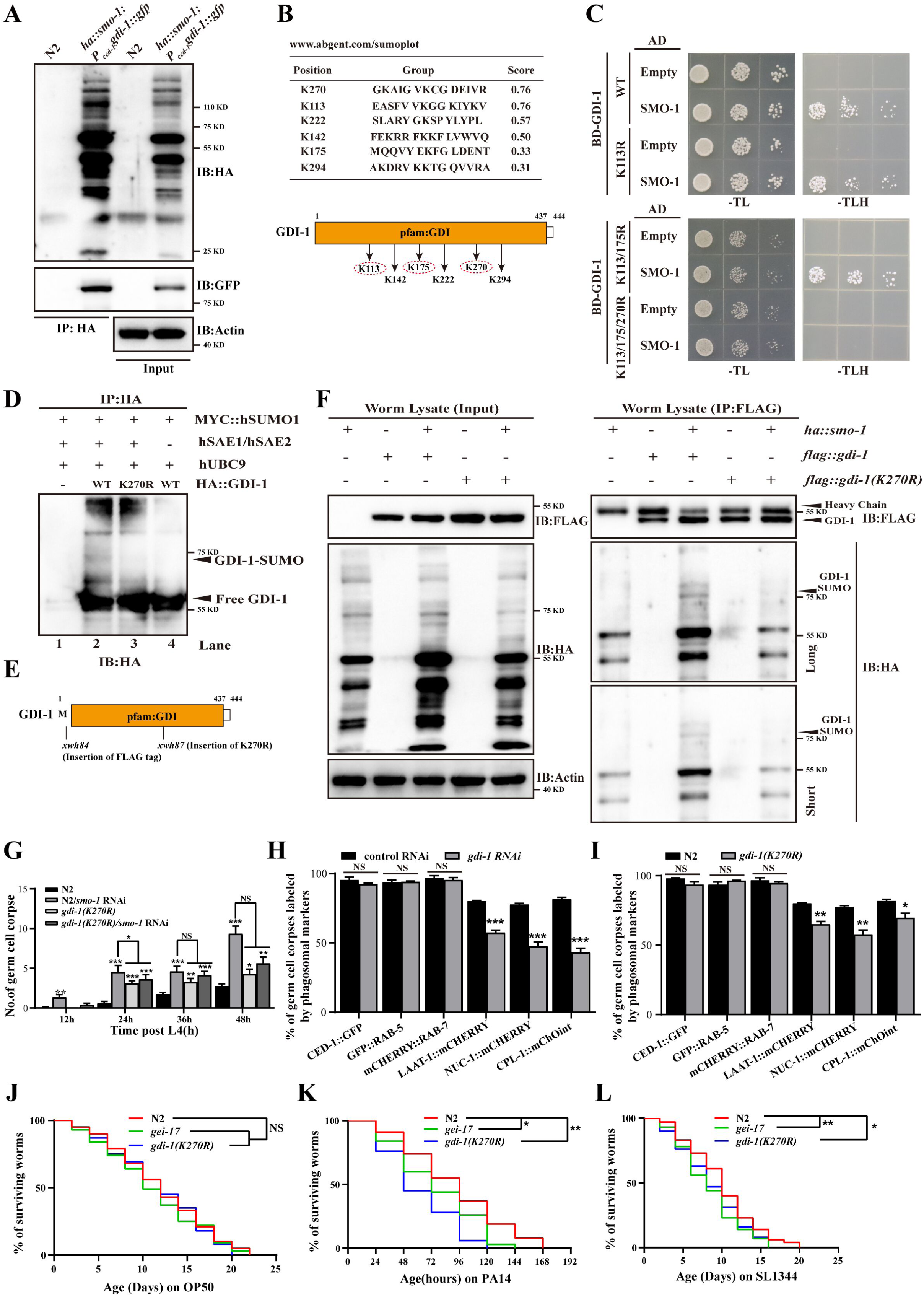
SUMOylation of GDI-1 at K270 is required for apoptotic cell degradation. (A) HA IP was performed on *ha::smo-1; Pced-1gdi-1::gfp* worms, followed by detection of the peptide of GDI-1 that interacts with SMO-1 by LC-MS/MS. (B) Predicted SUMOylation sites of GDI-1 using the SUMOplot web service (https://www.abcepta.com/sumoplot) (top), schematic illustration of the SUMOylation sites of GDI-1 (bottom). (C) Yeast two-hybrid assay of the interactions between GDI-1 (WT, K113R, K113R/175R, and K113/175/270R) and SMO-1. (D) *In vitro* SUMOylation of the wild-type and 270 lysine-to-arginine (K270R) mutant forms of GDI-1. The sumoylated-GDI-1 band is indicated by an arrowhead. SUMOylation of GDI-1(K270R) was not observed (lane 3 was compared with lanes 1, 2, and 4). (E) Schematic illustration of the mutation and tag insertions generated by CRISPR-Cas9 editing of the endogenous *gdi-1* loci. Amino acids near insertion or mutation sites are indicated. (F) SUMOylation of GDI-1 was examined in worms carrying *ha::smo-1* and *flag::gdi-1* (WT and K270R). Long and short designate long and short exposure times, respectively. (G) The different stages (h post L4) of germ cell corpses were quantified (mean ± SEM) in the indicated strains. Fifteen adult worms were scored at each stage for each strain. (H and I) Cell corpses positive for phagosome markers in control/*gdi-1* RNAi (H) and N2, gdi-1(K270R) (I) were quantified. A total of ≥100 cell corpses were analyzed for each marker, and data were derived from three replicates. (J-L) OP50, PA14, and SL1344 survival assays in N2, *gei-17*, and *gdi-1(K270R)* animals. Three biological replicates were used with 100 worms per replicate. The unpaired *t* test was performed in this figure. *P < 0.05, **P < 0.01, ***P < 0.001, NS, no significance. All error bars indicate the mean ± SEM.

**Table 2.** The peptides identified for GDI-1 in LC-MS/MS in *C. elegans*.

| Sequencec | Start | End | Score | Intensity |
| --- | --- | --- | --- | --- |
| FGLDENTADFTGHALALYR | 176 | 194 | 542.44 | 4.40E+09 |
| FGLDENTADFTGHALALYRDDEHK | 176 | 199 | 274.8 | 1.40E+09 |
| NNYYGGESASLTPLEQLYEK | 36 | 55 | 216.27 | 2.10E+10 |
| VPADEMEALATSLMGMFEK | 120 | 138 | 214.45 | 3.70E+09 |
| DDEHKNQPYAPAVEK | 195 | 209 | 203.7 | 2.00E+09 |
| EDTWQGLDPHNSTMQQVYEK | 156 | 175 | 189.1 | 4.40E+08 |
| FLVWVQQFDENK | 144 | 155 | 161.76 | 3.50E+08 |
| PVDEIVMENGK | 255 | 265 | 159.79 | 1.30E+07 |
| LSAIYGGTYMLDKPVDEIVMENGK | 242 | 265 | 156.25 | 2.30E+09 |
| CGDEIVR | 271 | 277 | 155.73 | 4.30E+09 |
| IRLYSDSLAR | 210 | 219 | 150.1 | 4.20E+07 |
| HYDIYISCCSNTNMVTPK | 330 | 347 | 145.29 | 7.70E+07 |
| VPADEMEALATSLMGMFEKR | 120 | 139 | 134.29 | 1.40E+08 |
| FIQISDVYEPSDLGSESQIFISQSYDATT<br>HFETTCK | 381 | 416 | 131.97 | 1.50E+08 |
| LYSDSLAR | 212 | 219 | 130.77 | 1.70E+10 |
| NNYYGGESASLTPLEQLYEKFHGPQAK | 36 | 62 | 129.23 | 7.80E+07 |
| DVLNMFER | 417 | 424 | 126.67 | 1.90E+10 |
| LSAIYGGTYMLDK | 242 | 254 | 121.08 | 4.50E+07 |
| FHGPQAKPQQEMGR | 56 | 69 | 121.02 | 2.10E+09 |
| PQQEMGR | 63 | 69 | 120.9 | 5.10E+09 |
| VLHIDRNNYYGGESASLTPLEQLYEK | 30 | 55 | 118.9 | 1.60E+09 |
| GRDWNVDLIPK | 70 | 80 | 118.71 | 1.00E+10 |
| AICLLNHPIPTNDAQSCQIIPQK | 301 | 325 | 118.22 | 1.10E+08 |
| IYKVPADEMEALATSLMGMFEK | 117 | 138 | 116.45 | 2.50E+09 |
| DWNVDLIPK | 72 | 80 | 104.2 | 6.50E+09 |
| ECIISGMLSVSGK | 16 | 28 | 103.34 | 1.40E+09 |
| NQPYAPAVEK | 200 | 209 | 100.84 | 9.90E+09 |
| AIGVKCGDEIVR | 266 | 277 | 99.752 | 9.00E+07 |
| FLMANGPLVK | 81 | 90 | 98.523 | 4.50E+09 |
| EFDFTNITHLSLNDQE | 429 | 444 | 94.538 | 3.00E+07 |
| FLVWVQQFDENKEDTWQGLDPHNST<br>MQQVYEK | 144 | 175 | 92.609 | 2.20E+08 |
| QIYCDPSYAK | 280 | 289 | 89.247 | 6.50E+09 |
| SIEASFVVK | 105 | 113 | 71.501 | 6.80E+09 |
| QIYCDPSYAKDR | 280 | 291 | 67.519 | 1.60E+08 |
| NQPYAPAVEKIR | 200 | 211 | 56.951 | 4.50E+07 |
| DDEHKNQPYAPAVEKIR | 195 | 211 | 53.453 | 1.40E+08 |
| GKQIYCDPSYAK | 278 | 289 | 28.433 | 2.70E+07 |
| LSAIYGGTYMLDKPVDEIVMENGKAIG<br>VK | 242 | 270 | 24.612 | 1.70E+07 |

To verify K270 as the SUMO modification site on GDI-1, we performed *in vitro* SUMOylation reactions with recombinant proteins expressed and purified from *E. coli*. We found that SUMO modification was indeed abolished in the GDI-1(K270R) variant (Fig. 4D), thus supporting K270 as the likely SUMOylation site. We then generated the *xwh87(flag::gdi-1(K270R))* worm strain by CRISPR-Cas9 editing that expressed a flag-tagged *gdi-1* that had a K→R conversion at the SUMOylation site (Fig. 4E). Western blot assays of IP lysates revealed that the high molecular weight species were absent in the *xwh87* strain (Fig. 4F), thus demonstrating K270 as the site required for GDI-1 SUMOylation.

To further explore the role of GDI-1 SUMOylation in apoptotic cell clearance, CRISPR-Cas9 was used to introduce the K270R mutation at the endogenous *gdi-1* locus, resulting in the *xwh89(gdi-1(K270R))* mutant strain, which exhibited significantly increased numbers of germline cell corpses compared to that in N2 worms (Fig. 4G). Moreover, RNAi silencing of *smo-1* in these *gdi-1(K270R)* mutants did not significantly affect germ cell corpse number (Fig. 4G), supporting K270 as the GDI-1 SUMOylation site necessary to activate apoptotic cell clearance. Given our above results showing that SUMOylation positively regulates phagosomal recruitment of LAAT-1 and acidification of the phagosome (Fig. 2G), we next examined SUMO modification of GDI-1 was required for recruitment of phagosomal maturation factors. Fluorescence microscopy revealed both *gdi-1* RNAi knockdown worms and *gdi-1(K270R)* mutants expressing LAAT-1::GFP, NUC-1::mCHERRY, or CPL-1::mChoint had significantly lower proportion of germline cell corpses with these markers around phagosomes compared to WT (Fig. 4H,I), which was consistent with our above results in *smo-1* knockdown worms. These data indicated that SUMOylation of GDI-1 residue K270 promotes apoptotic cell clearance by influencing phagosome maturation *in vivo*.

We then investigated whether endosomal/lysosomal degradation activity was affected in *gei-17* mutants and *gdi-1(K270R)* mutants by introducing the *arIs36(P_heat-shock_ssGFP)* expression vector into these strains. Quantification of ssGFP uptake and degradation in coelomocytes over a 36h time course showed that ssGFP began to accumulate in the body cavity (pseudocoelom) of *arIs36* worms by one hour post-heat shock (hphs), and was subsequently transported to coelomocytes (*Figure S4A*). The ssGFP signal progressively decreased in the body cavity by 12 hphs, conjointly with increased ssGFP GFP signal in coelomocytes, and was no longer detectable in the body cavity or coelomocytes by 24 hphs (Fig. EV4A), suggesting its complete degradation by coelomocytes. Disruption of SUMO modification did not obviously affect initial ssGFP uptake into coelomocytes, but ssGFP could still be detected in both the body cavity and coelomocytes of *gei-17* mutants or *gdi-1(K270R)* mutants at 36 hphs (Fig. EV4A), suggesting that loss of GDI-1 SUMOylation led to impaired endosomal/lysosomal activity.

To directly observe the process of cell corpse degradation, we introduced the *ujIs113*(*P_pie-1_H2B::mCherry; P_nhr-2_HIS-24::mCherry*) vector construct into WT and *gdi-1(K270R)* worms. Detection of mCherry signal by fluorescence microscopy showed condensed chromatin in germ cell corpses within early stage phagosomes and disappeared within ∼30 minutes in WT worms. By contrast, mCHERRY signal persisted for longer than 90 min in the *ujIs113; gdi-1(K270R)* strain (Fig. EV4B and C), suggesting that chromatin degradation in cell corpses was substantially delayed in the absence of GDI-1 SUMOylation. These data cumulative demonstrated that cell corpse accumulation in *gdi-1(K270R)* animals was due to defects in lysosomal degradation of engulfed cell corpses.

Although lacking innate immunity, the *C. elegans* immune system to some extent resembles the human innate immune system and can respond to pathogen infection by upregulating antimicrobial peptide expression. As apoptotic cell clearance has been linked to innate immune function in *C. elegans,* we also examined whether GDI-1 SUMOylation contributed to regulating innate immunity by analyzing the lifespan of WT N2, *gei-17* knockout, and *gdi-1(K270R)* worms fed on OP50, *P. aeruginosa* PA14, and *S. typhimurium* SL1344, respectively. Although mutants fed with OP50, the standard nematode diet, exhibited similar lifespan to that of N2, both *gei-17* and *gdi-1(K270R)* strains became infected and showed significantly reduced lifespans upon feeding with the pathogenic PA14 and SL1344 bacterial strains (Figs. 4J-L and EV4D-F), suggesting that GDI-1 SUMOylation at K270 was required for *C. elegans* resistance to bacterial pathogens.

### GDI-1 regulates RAB-1 activity during cell corpse clearance

To better define possible connections between GDI-1 SUMOylation and its function in apoptotic cell clearance, we investigated whether SUMO modification altered the protein stability, protein–protein interactions, subcellular localization, or catalytic activity of GDI-1. Western blots showed that endogenous GDI-1 protein levels did not significantly differ between the N2, *gei-17,* and *gdi-1(K270R)* strains (Fig. 5A), suggesting that SUMOylation does not affect GDI-1 stability. As GDI family proteins regulate GDP/GTP exchange of RAB proteins by stabilizing membrane-associated, prenylated RABs in the cytosol, or restoring them to the membrane for activation, we next sought to identify the specific RAB binding partner of GDI-1 involved in apoptotic cell clearance. To this end, we individually expressed 31 RAB proteins from *C. elegans* (Gallegos, Balakrishnan et al., 2012) in Y2H assays and found that GDI-1 could interact with RAB-1, RAB-28, and RAB-Y1 (Figs. 5B and EV5A-F).

**Figure 5.**
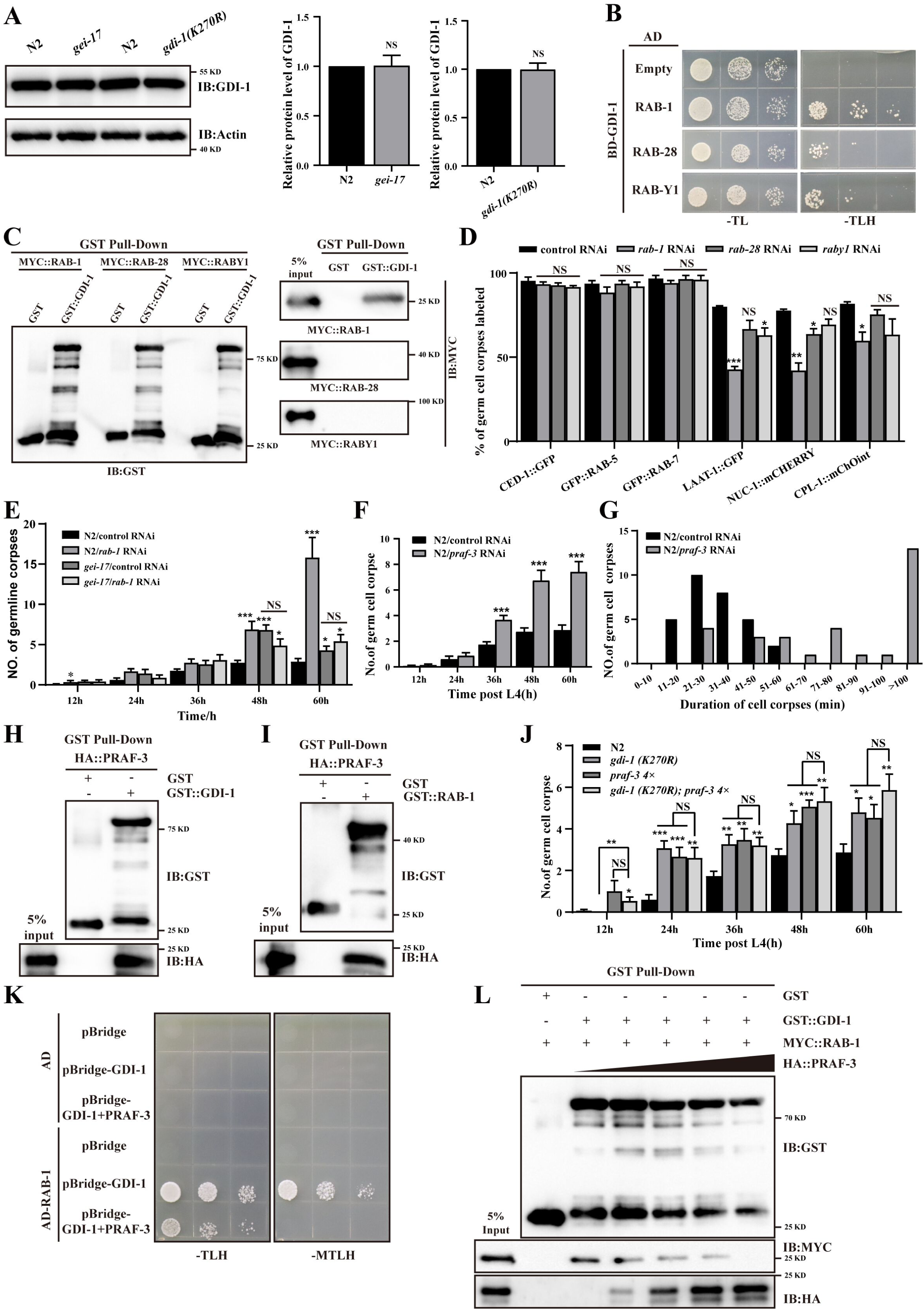
GDI-1 regulates RAB-1 activity during cell corpse clearance. (A) Endogenous GDI-1 protein levels were examined by immunoblot analysis in N2, *gei-17*, and *gdi-1(K270R)* worms. Graphs show quantification of the protein level of GDI-1 in the indicate strains. (B) The yeast two-hybrid assay of the interaction between GDI-1 and RAB-1/RAB-28/RAB-Y1. (C) Germ cell corpses positive for phagosome markers in N2 treated with control or *rab-1/rab-28/rab-y1* RNAi were quantified. A total of ≥100 cell corpses were analyzed for each marker; and data were derived from three replicates. (D, H, and I) The interactions between GDI-1 and RAB-1/RAB-28/RAB-Y1 (D), PRAF-3 and RAB-1 (H), PRAF-3 and GDI-1 (I) were examined by GST pull-down assays. (E, F, and J) The different stages (h post L4) of germ cell corpses were quantified (mean ± SEM) in the indicated strains. Fifteen adult worms were scored at each stage for each strain. (G) Four-dimensional microscopy analysis of cell corpse duration was performed in N2/control and N2/*praf-3* RNAi worms. The persistence of 30 cell corpses from the embryos was monitored. (K) The yeast three-hybrid assay of GDI-1 and PRAF-3 competitively combined with RAB-1. Constitutive Gal4-BD GDI-1 expression and inducible PRAF-3 expression elements were cloned into pBridge. PRAF-3 expression was induced when methionine was dropout. -TLH, medium lacking Trp, Leu, and His; -MTLH, medium lacking Met, Trp, Leu, and His. (L) GDI-1 and PRAF-3 competitively combined with RAB-1 was examined by GST pulldown assays. Different concentrations of PRAF-3 were added to the system, in which GDI-1 interacted with RAB-1, and RAB-1 and PRAF-3 were detected under the pull-down of GDI-1 using IB. From left to right lanes, PRAF-3 was added in amounts of 0, 1, 2, 3, 4, and 5 µg. The unpaired *t* test was performed in this figure. *P < 0.05, **P < 0.01, ***P < 0.001, NS, no significance. All error bars indicate the mean ± SEM.

Subsequent co-IP assays in HEK 293T cells expressing each respective candidate revealed that RAB-1 and RAB-Y1 could interact with GDI-1 (Fig. EV5G-I). GST pull-down assays showed that only recombinant RAB-1 protein could be pulled down by GST-GDI-1, and not by GST alone (Fig. 5C), indicating that RAB-1 directly interacted with GDI-1. In addition, fluorescence microscopy showed that germline cell corpses of *rab-1* RNAi knockdown worms expressing LAAT-1::GFP, NUC-1::mCHERRY, or CPL-1::mChoint had significantly decreased signal from these markers around phagosomes (Fig. 5D), similar to the results of *gdi-1* knockdown (Fig. 4H). In addition, *rab-1* knockdown in N2 and *gei-17* showed that apoptotic germ cell abundance significantly increased in *gei-17* mutants with *rab-1* knockdown compared to *gei-17* and N2 worms with or without *rab-1* RNAi treatment (Fig. 5E), suggesting that *gei-17* and *rab-1* functioned in the same genetic pathway. These results collectively suggested that GDI-1 SUMOylation could promote phagosome maturation in apoptotic cell clearance via interaction with RAB-1.

As membrane-bound GDI displacement factors (GDFs) mediate RAB-GDP dissociation from the RAB-GDP-GDI complex to catalyze Rab insertion into cognate membranes, we therefore tested whether PRAF-3, the only *C. elegans* ortholog of mammalian GDFs, also participated in apoptotic cell clearance. We found that RNAi suppression of *praf-3* led to an age-dependent increase in cell corpse numbers in germline cells compared to that in vector controls (Fig. 5F). To test whether *praf-3* knockdown increased cell corpse number through excessive apoptosis or defective phagosomal activity, we quantified cell corpses under *praf-3* knockdown in transgenic *P_ced-1_rde-1; rde-1* worms (with RNAi suppression of phagosome formation) and *P_egl-1_rde-1; rde-1* worms (with RNAi suppression of apoptosis). We observed *praf-3* knockdown led to increased cell corpse abundance in *P_ced-1_rde-1; rde-1,* but not *P_egl-1_rde-1; rde-1* worms (Fig. EV5J,K), suggesting that *praf-3* functioned in cell corpse clearance rather than induction of apoptosis. In addition, four-dimensional microcopy analysis of the timespan for germ cell corpse persistence showed that germ cell corpse degradation required an average of 36.5±2.8 min in WT animals, whereas cell corpse degradation required 112.8±4.1 min in *praf-3* knockdown gonads (Fig. 5G).

Subsequent GST pull-downs showed that recombinant HA::PRAF-3 could be immunoprecipitated with GST-GDI-1, but not with GST alone (Fig. 5H), indicating that PRAF-3 could directly interact with GDI-1 *in vitro*. Furthermore, GST pull-downs and co-IP assays with RAB-1 showed that HA::PRAF-3 could also be immunoprecipitated by GST::RAB-1 or MYC::RAB-1, respectively (Figs. 5I and EV5L). As PRAF-3, GDI-1 and RAB-1 appeared to directly interact with each other, we also examined whether GDI-1 competed with PRAF-3 for binding with RAB-1 using Yeast 3-Hybrid (Y3H) and GST pull-down assays. We found that PRAF-3 could disrupt GDI-1 interaction with RAB-1 (Fig. 5K) in Y3H assays. Similarly, adding increasing concentrations of PRAF-3 to GDI-1 incubations with RAB-1 resulted in a dose-dependent decrease in RAB-1–GDI-1 interactions (Fig. 5L). These data suggested that PRAF-3 could compete with GDI-1 to bind RAB-1 to modulate apoptotic cell clearance in *C. elegans*.

### GDI-1 SUMOylation regulates RAB-1-mediated vesicular transport of RAB-7 from ER to Golgi

Given the above direct GDI-1-RAB-1 interactions, we next investigated the function of GDI-1 SUMOylation in regulating its interactions with RAB-1. First, WB analysis of RAB-1 protein contents in *gdi-1(K270R)* mutants showed that total RAB-1 levels did not significantly differ from that in N2 (Fig. 6A,B). However, isolation of the plasma membrane and cytoplasmic protein fractions in N2, *gdi-1(K270R)*, and *praf-3(xwh92)* mutants followed by WB detection of endogenous RAB-1 revealed that RAB-1 levels were significantly increased in the cytoplasmic fraction and concomitantly decreased in the membrane fractions from *gdi-1(K270R)* and *praf-3* mutants (Fig. 6C-E). These results suggested that SUMO modification of GDI-1 affected the intracellular localization of RAB-1. However, GST pull-down assays showed no significant difference in recombinant MYC::RAB-1 levels pulled down by GST-GDI-1 versus GST-GDI-1(K270R) (Fig. EV6A,B). As *in vitro* GST pull-downs might not accurately reflect the function of SUMOylation in regulating protein-protein interactions *in vivo,* we therefore conducted co-IP assays in *gdi-1(K270R)* and *gei-17(xwh81)* mutants with impaired GDI-1 SUMOylation. Subsequent WB detection of RAB-1 co-IP extracts of GD-1 showed higher levels of DI-1-RAB-1 interaction in the *gdi-1(K270R)* and *gei-17(xwh81)* mutants compared to that in N2 (Fig. 6F,G). Similar results were obtained with Flag-tagged GDI-1 in *flag::gdi-1(K270R)* and *gei-17; flag-gdi-1* worm lysates (Fig. EV6C,D). Based on these results, we hypothesized that unSUMOylated GDI-1 may undergo stronger interactions with GDP-bound RAB-1 *in vivo*, maintaining RAB-1 in the inactive state.

**Figure 6.**
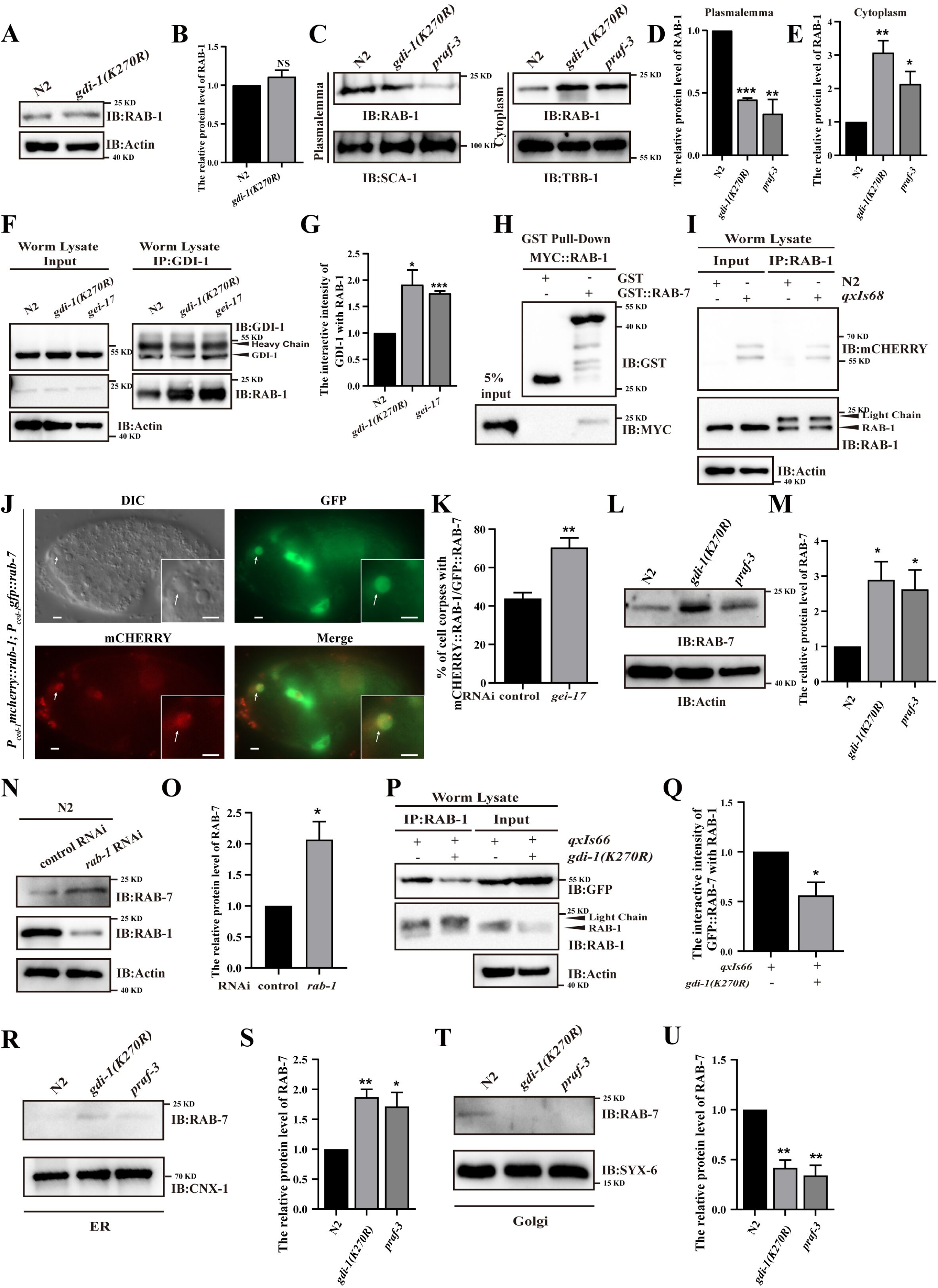
GDI-1 SUMOylation regulates RAB-1-mediated vesicular transport of RAB-7 from ER to Golgi. (A) Endogenous RAB-1 protein levels were examined by immunoblotting in N2 and *gdi-1(K270R)* worms. (B) Graphs showing the quantification of the protein level of GDI-1 in N2 and *gdi-1(K270R)* worms. (C) Endogenous RAB-1 protein levels were examined by immunoblot analysis in N2, *gdi-1(K270R)*, and *praf-3* plasmalemma or cytoplasm. SCA-1 is a plasmalemma marker and TBB-1 is a cytoplasmic marker. (D and E) The graphs show quantification of the protein level of RAB-1 in the plasmalemma (D) and cytoplasm (E) using ImageJ software. (F) GDI-1 IP was performed, followed by detection of the interaction intensity between GDI-1 and RAB-1 in N2, *gei-17*, and *gdi-1(K270R)* worms with an anti-RAB-1 antibody. (G) The graph shows the protein level of RAB-1/GDI-1. The ratio of RAB-1 to GDI-1 was determined and normalized to one-fold in N2. (H) GST pull-down assay of the interaction between RAB-1 and RAB-7. (I) RAB-1 IP was performed, followed by detection of the interaction between RAB-1 and RAB-7 in N2, *qxIs68 (P_ced-1_mcherry::rab-7)* worms with anti-mCHERRY antibody. (J) Colocalization of GFP::RAB-7 and mCHERRY::RAB-1 in N2 embryos. Arrows indicate cell corpses. The boxed regions are magnified (2×) in the inset. Bars, 2 µm. (K) Quantification graph of GFP::RAB-7 and mCHERRY::RAB-1 colocalization on cell corpses in control and *gei-17* RNAi embryos. Percentage refers to the ratio of mCHERRY::RAB-1 to GFP::RAB-7. (L and N) Endogenous RAB-7 protein levels were examined by immunoblotting in N2/*gdi-1(K270R)*/*praf-3* (L) and control/*rab-1* RNAi (N) worms. (M and O) Graphs show the quantification of the protein level of RAB-7 in N2/*gdi-1(K270R)*/*praf-3* (M) and control/*rab-1* RNAi (O) worms. (P) RAB-1 IP was performed, followed by the detection of the interaction intensity between RAB-1 and RAB-7 in *qxIs66 (P_ced-1_gfp::rab-7)* and *gdi-1(K270R); qxIs66 (P_ced-1_gfp::rab-7)* worms with an anti-GFP antibody. (Q) The graph shows protein levels of GFP::RAB-7/RAB-1. The ratio of GFP::RAB-7 to RAB-1 was determined and normalized one-fold in *qxIs66*. (R and T) Endogenous RAB-7 protein levels were examined by immunoblot analysis in N2, *gdi-1(K270R)*, and *praf-3* in endoplasmic reticulum (ER) (R) or Golgi apparatus (T). CNX-1 is an ER marker and SYX-6 is a Golgi marker. (S and U) The graphs show quantification of the protein level of RAB-7 in the ER (S) and Golgi apparatus (U) using ImageJ software. The unpaired *t* test was performed in this figure. *P < 0.05, **P < 0.01, ***P < 0.001. All error bars indicate the mean ± SEM.

As RAB1 can reportedly interact with RAB7 in *Drosophila* (Guruharsha, Rual et al., 2011), we also tested whether RAB-1 could interact with RAB-7 in *C. elegans*. We found that GST-RAB-7, but not GST, could pull down recombinant MYC::RAB-1 *in vitro*, indicating that RAB-7 could directly interact with RAB-1 (Fig. 6H). In addition, co-IP assays in the transgenic *qxIs68(P_ced-1_mcherry::rab-7)* strain, expressing an mCHERRY::RAB-7 fusion reporter driven by the *ced-1* promoter, showed that RAB-1 could be co-precipitated with RAB-7 from crude extracts using mCHERRY antibodies (Fig. 6I). Fluorescence microscopy further revealed that mCHERRY::RAB-1 co-localized with a GFP::RAB-7 reporter on the surface of cell corpses in *C. elegans* (Fig. 6J), and this co-localization increased in worms with *gei-17* knockdown (Fig. 6K). Furthermore, co-IP assays in HEK 293T cells also indicated that RAB-1 could interact with RAB-7 (Fig. EV6E). These results demonstrated that RAB-7 could indeed interact with RAB-1 in *C. elegans*.

To further investigate the mechanism through which SUMO modification of GDI-1 might regulate RAB-1 interaction with RAB-7, we performed GST pull-down assays and found that HA::RAB-7 protein could not be pulled down by GST-GDI-1 (Fig. EV6F), indicating that RAB-7 could not directly interact with GDI-1. Interestingly, endogenous RAB-7 levels were significantly increased in *gdi-1(K270R)*, *praf-3(xwh92)*, and *rab-1* knockdown worms compared to that in controls (Fig. 6L-O). Moreover, exogenous GFP::RAB-7 expression was also significantly increased in both *gdi-1(K270R); P_ced-1_gfp::rab-7* and *rab-1* RNAi-treated *P_ced-1_gfp::rab-7* worms (Fig. EV6G-J). WB analysis of RAB-7 contents immunoprecipitated by anti-RAB-1 antibody indicated that RAB-1 interaction with RAB-7 was reduced in *qxIs66; gdi-1 (K270R)* (Fig. 6P,Q), which might be due to stronger GDI-1 interaction with RAB-1 in the cytoplasm where it lacks SUMO modification, consequently blocking RAB-1 binding with RAB-7. Fluorescence microscopy showed no significant difference in the percentage of mCHERRY::RAB-1 signal localized to the surface of cell corpses between *P_ced-1_mcherry::rab-1; gdi-1 (K270R)* and *P_ced-1_mcherry::rab-1* worms (Fig. EV6K), whereas the percentage of mCHERRY::RAB-1 on the cell corpse surface was significantly lower in *rab-7* RNAi-treated *P_ced-1_mcherry::rab-1* animals compared with *P_ced-1_mcherry::rab-1* controls (Fig. EV6L). These results suggested that RAB-7 may function downstream of RAB-1 activation in cell corpse clearance.

As RAB-7 is known to mediate lysosome recruitment and subsequent fusion to with phagosomes in phagolysosome formation, we therefore investigated whether RAB-7 was degraded via the endo-lysosomal pathway. To this end, we blocked lysosomal acidification required for its enzymatic function by pretreating *P_ced-1_gfp:: rab-7* worms with chloroquine (Chl). Western blots indicated that endogenous RAB-7 contents significantly increased compared to vehicle controls (Fig. EV6M,N). As the ubiquitin-proteasome pathway is the other major protein degradation pathway in eukaryotic cells, we also chemically inhibited this pathway with MG-132 and observed a similar increase in RAB-7 accumulation (Fig. EV6M,N). These results suggested that RAB-7 levels could be simultaneously modulated through both lysosome-mediated and ubiquitin-proteasome-mediated degradation. We then pretreated *gdi-1(K270R)*; *P_ced-1_gfp::rab-7* worms with Chl and MG-132, individually, and found that endogenous RAB-7 accumulated to high levels in the MG-132, but not in the Chl group (Fig. EV6O,P), indicating that the increase in RAB-7 contents resulting from impaired GDI-1 SUMOylation occurred independently of the proteasomal degradation pathway.

As mammalian Rab1A and Rab1B are predominantly found on endoplasmic reticulum (ER) membrane and Golgi apparatus and function redundantly in vesicular transport from ER to Golgi, we next examined whether RAB-1 regulates vesicular transport of RAB-7 between the ER and successive Golgi compartments by isolating the ER and Golgi protein fractions from N2, *gdi-1(K270R)*, and *praf-3(xwh92)* mutants. WB analysis showed that endogenous RAB-7 levels significantly increased in ER with a concomitant decrease in Golgi of *gdi-1(K270R)* and *praf-3* mutants (Fig. 6R-U), suggesting that SUMO modification of GDI-1 affects RAB-7 localization. Additionally, we overexpressed recombinant mCHERRY::RAB7A in HEK293T cells with concurrent *rab1a* knockdown by shRNA. Upon extracting ER and Golgi, WB detection of mCHERRY indicated that RAB7A protein levels significantly increased in ER, and again showed a concomitant decrease in Golgi under *rab1a* knockdown in HEK293T cells (Fig. EV6R-U). These cumulative results demonstrated that RAB1 mediates the vesicular transport of RAB-7 from ER to Golgi to promote apoptotic cell clearance.

### SUMOylation of GDI1 modulates efferocytosis in mammals

Given the above evidence that GDI-1 SUMOylation by GEI-17 is required for its regulation of RAB-1 in *C. elegans*, we next investigated whether these regulatory interactions also occur in mammals. Co-IP assays in HEK293T cells co-expressing an HA tagged human homolog of GDI-1, HA::hGDI1, a MYC-tagged human homolog of GEI-17, MYC::PIAS1, and recombinant human MYC::RAB1A, indicated that HA::hGDI-1 could interact with both recombinant MYC::PIAS1 and MYC::RAB1A (Fig. 7A,B). Subsequent GST pull-down assays showed that the recombinant MYC::PIAS1 or MYC::RAB1A proteins could be pulled down by GST-hGDI1, but not by GST alone (Fig. EV7A,B), indicating that hGDI1 could directly interact with both PIAS1 and RAB1A *in vitro*. Interestingly, co-IP assays in HEK293T cells showed that hGDI1 could interact with *C. elegans* proteins, GEI-17 and RAB-1, while GDI-1 could interact with their human homologs, PIAS1 and RAB1A (Fig. 7C-F), suggesting that these interactions were likely conserved between worms and humans.

**Figure 7.**
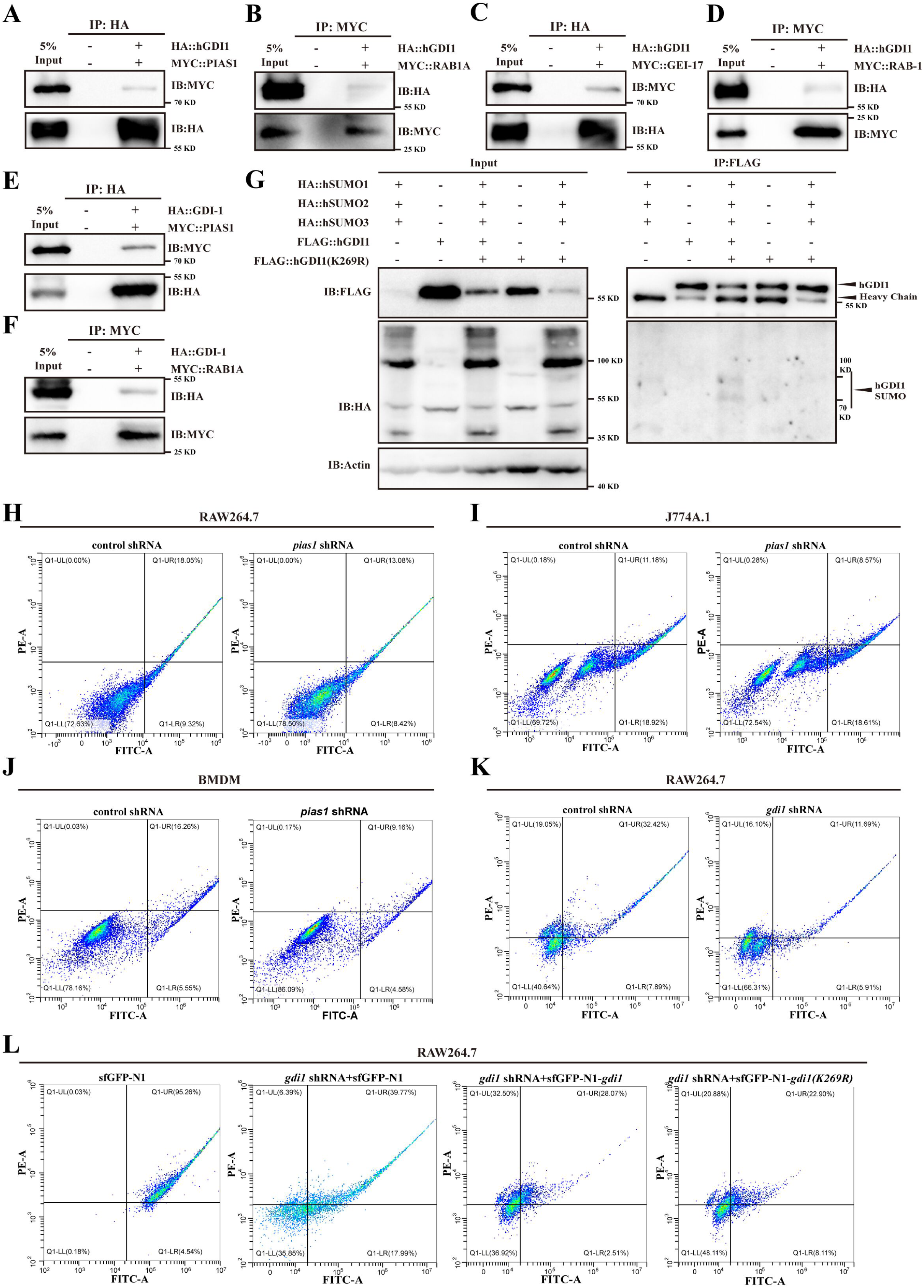
SUMOylation of GDI1 modulates efferocytosis in mammals. (A-F) The interactions between hGDI1 and PIAS1 (A), hGDI1 and RAB1A (B), hGDI1 and GEI-17 (C), hGDI1 and RAB-1 (D), GDI-1 and PIAS1 (E), and GDI-1 and RAB1A (F) were examined by CO-IP in HEK293T cells. (G) SUMOylation of the wild-type and 269 lysine-to-arginine (K270R) mutant forms of hGDI1. The SUMOylated-hGDI1 band is indicated by an arrowhead. (H-J) Phagocytosis rates of control or *pias1* shRNA-treated RAW264.7 (H), J774A.1 (I), and BMDM (J) phagocytes were analyzed using flow cytometry. (K) Flow cytometry phagocytosis assays of control or *gdi1* shRNA-treated RAW264.7. (L) Phagocytosis rates of sfGFP-N1, *gdi1* shRNA+sfGFP-N1, *gdi1* shRNA+sfGFP-N1-*gdi1*, and *gdi1* shRNA+sfGFP-N1-*gdi1(K269R)* treated RAW264.7 phagocytes were analyzed using flow cytometry.

The above results indicate that the interactions of GDI-1 with GEI-17 and RAB-1 were evolutionarily conserved. Subsequently, we investigated whether mammalian GDI1 was modified by SUMOylation and whether the SUMOylation site of GDI-1 in *C. elegans* was conserved. Sequence comparison revealed that the amino acid sequence of GDI-1 is conserved in *C. elegans*, mice, and humans, where the lysine at position 270 of GDI-1 corresponds to the amino acid at position 269 of murine and human proteins (Fig. EV7C). Moreover, FLAG IP assays in HEK293T cells expressing HA::hSUMO1/2/3 and FLAG::hGDI1(WT and K269R) showed that hGDI1 was SUMOylated, whereas hGDI1(K269R) was not (Fig. 7G). These findings suggest that hGDI1 was SUMOylated via lysine at position 269. To investigate whether SUMO modifications also affected apoptotic cell clearance in mammals, we first tested the efficiency of shRNA (*pGreen-puro*) knockdown of *pias1* and *gdi1* in RAW264.7 macrophage cells by qPCR (Fig. EV7D,E). To perform efferocytosis assays, in addition to RAW264.7 cells, we also induced *pias1* knockdown with the same *pias1-*shRNA vector in J774A.1 and BMDM macrophages, then pre-labeled Jurkat T with CellTracker red before inducing apoptosis by UV radiation. The apoptotic cells Jurkat T cells were incubated with macrophages for 6h, and fluorescence image analysis was used to quantify the ratio of cell tracker-positive cells to total GFP-positive cells. This analysis revealed that efferocytosis was significantly reduced in RAW264.7, J774A.1, and BMDM cells with *pias1* knockdown compared to that in controls (Fig. EV7F-K). Flow cytometry further illustrated this significant decrease in efferocytosis efficiency under *pias1* knockdown (Fig. 7H-J). These results indicated that blocking PIAS1-mediated SUMOylation negatively affected the phagocytosis of apoptotic cells by macrophages. To elucidate the role of GDI SUMOylation in mammalian efferocytosis, we initially induced *gdi1* knockdown in RAW264.7, and found that efferocytosis was significantly reduced in RAW264.7 with *gdi1* knockdown compared to that in the control (Figs. 7K and EV7L,M). We then overexpressed GDI1(WT and K269R) in the *gdi1* knockdown background. Flow cytometric analysis revealed that while wild-type GDI1 successfully restored efferocytosis in *gdi1*-deficient cells, the K269R mutant failed to do so (Fig. 7L). Collectively, these results demonstrate that GDI1 SUMOylation plays a crucial role in the regulation of efferocytosis in mammals.

## Discussion

Ubiquitin and the ubiquitin-related modifier SUMO function in various aspects of the cellular stress response, such as targeting misfolded proteins for degradation, regulating transcription, and facilitating recovery from stress (Dikic & Schulman, 2023, Zheng & Shabek, 2017). Previous studies have demonstrated that ubiquitination modifications are crucial for apoptotic cell clearance (Liu, Li et al., 2018, Yuan et al., 2022a, Yuan, Li et al., 2022b). However, whether SUMO modifications are involved in apoptotic cell clearance remains unclear, although it has been established that SUMO modifications are implicated in apoptosis in mammals (Li et al., 2021). Here, we found that the SUMO precursor molecule SMO-1, the E1 activator AOS-1/UBA-2, the E2 conjugating enzyme UBC-9, and the E3 ligase homologs GEI-17/MMS-21 are involved in the clearance of physiological apoptotic cells rather than in excessive apoptosis by using *C. elegans,* the suitable model organism to dissect SUMO modification. SUMOylation has been predominantly investigated in the development as well as in DNA damage response in *C. elegans* (Fergin et al., 2022). To the best of our knowledge, we provide the first evidence that SUMO modification regulates apoptotic cell clearance in *C. elegans*. We found that the two E3 ligases, *mms-21* and *gei-17,* are most likely not in the same genetic pathway for apoptotic cell clearance. We focus here on the function of GEI-17 in apoptotic cell clearance. A further study with more focus on the function of *mms-21* in apoptotic cell clearance is therefore suggested.

Previous proteomic analyses of SUMOylated worm proteins have relied on *smo-1* transgenes that encode HIS-tagged SUMO fused to the FLAG epitope or to GFP (Becker et al., 2013, Drabikowski et al., 2018). Kim H et al inserted HIS10 sequences directly after the initiator methionine codon of *smo-1* without any other sequences, which is fully functional and can be utilized in conjunction with mutagenesis studies to identify specific SUMO acceptor lysines in substrate proteins (Kim, Ding et al., 2021a, Kim et al., 2021b). Employing this affinity-based approach and mass spectrometry, they identified hundreds of *C. elegans* proteins that were likely to be direct SUMO conjugates. In the present study, we used CRISPR-Cas9 to insert the HA tag directly after the initiator codon of *smo-1* and identified 188 potentially SUMO-modified proteins in *C. elegans* under normal development conditions through immunoprecipitation with an anti-HA antibody followed by LC-MS/MS analysis (Table 1). Our findings complement published work identifying SUMO-regulated targets in *C. elegans* and suggest that numerous SUMOylation substrates remain to be identified. We found GDI-1 interacts with SMO-1/SUMO, as well as GEI-17, suggesting that GDI-1 may recruit these factors directly. Utilizing affinity enrichment of SUMO and mutagenesis of candidate SUMO-acceptor sites, we determined that the lysine residue at position 270 is required for the SUMOylation of GDI-1. Mutation of lysine 270 to arginine in GDI-1 resulted in impaired degradation in apoptotic cells without altering GDI-1 protein content (Fig. 8).

**Figure 8.**
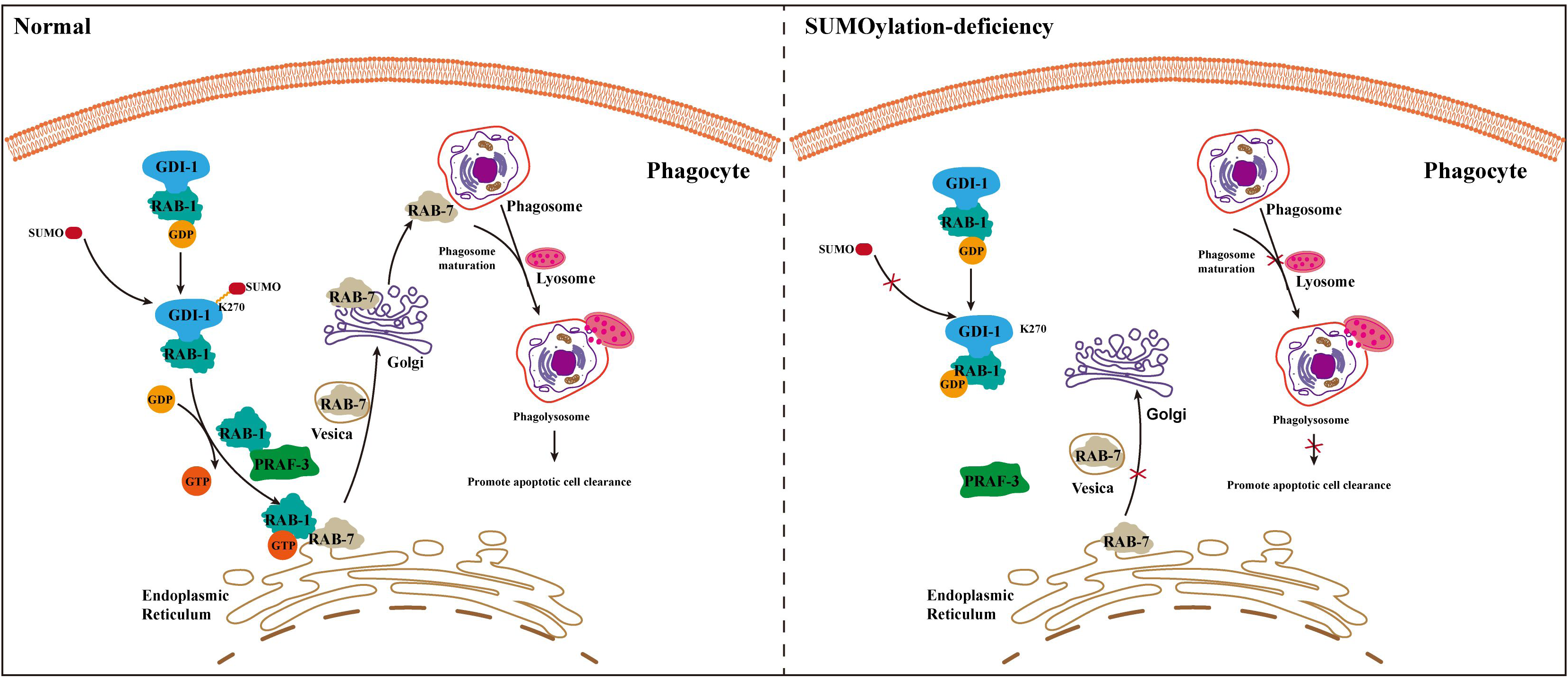
Schematic summary of SUMOylation of GDI-1 promotes apoptotic cell clearance. During engulfment, GDI-1-K270 is SUMOylated, SUMOylated GDI-1 promotes the release of RAB-1-GDP, and PRAF-3 converts RAB-1-GDP to RAB-1-GTP. The active form of RAB-1 aids in the vesicular transport of RAB-7 from the endoplasmic reticulum to the Golgi, and ultimately RAB-7 localizes to the phagosome, promoting phagosome maturation and recruitment of lysosomes, which in turn promotes phagosomal degradation. In the absence of SUMOylation, GDI-1 is unable to SUMOylate, leaving RAB-1 in its GDP form, and RAB-7 cannot be transported from the endoplasmic reticulum to the Golgi, excessively accumulating in the endoplasmic reticulum, and consequently suppressing phagosomal degradation.

GDI binds prenylated GDP-conjugated Rabs in the cytosol or membrane, mediating the transport of GDP-Rabs or their recovery from the target membrane (Bar, Charar et al., 2018, Gilbert & Burd, 2001, Muller & Goody, 2018). The Rab family of proteins is a crucial regulator of membrane trafficking and plays a significant role in numerous cellular processes, including phagocytosis. Research has demonstrated that RAB-5 regulates the fusion of early endosomes, RAB-2 and RAB-14 regulate the maturation of phagosomes in parallel, and RAB-7 mediates the maturation of early endosomes to late endosomes and the recruitment and fusion of lysosomes during the degradation of apoptotic cells in *C. elegans* (Mangahas, Yu et al., 2008, Zhang, Jiang et al., 2022). Our investigation revealed that GDI-1 interacts with RAB-1 and that knockdown of RAB-1 results in impaired phagosome recruitment to the lysosome, suggesting that RAB-1 and SUMO modification function in the same genetic pathway for apoptotic cell clearance. PRAF-3 (GDI displacement factor GDF in *C. elegans*) interacts with both GDI-1 and RAB-1, and GDI-1 binds RAB-1 competitively with PRAF-3. Mutations in PRAF-3 affect the clearance of apoptotic cells. Notably, we demonstrated that PRAF-3 and SUMO modification of GDI-1 function in the same genetic pathway for the clearance of apoptotic cells in *C. elegans* (Fig. 8).

RAB1 is located at the endoplasmic reticulum exit site (ERES) and pre-Golgi intermediate (pre-GIC), and mediates endoplasmic reticulum-Golgi transport (Zhang et al., 2022). Here, we found that SUMOylation of GDI-1 affects the localization of RAB-1, increasing its cytoplasmic localization and decreasing its membrane localization, and that impaired SUMOylation of GDI-1 leads to GDI-1 binding to more RAB-1. SUMOylation of GDI-1 appears to aid in the release of the GDP form of RAB-1, facilitating its membrane localization in its active form. *Drosophila* protein interaction mapping has revealed a possible interaction between RAB1 and RAB7 (Guruharsha et al., 2011). Previous research has demonstrated that SUMOylated Rab7 is recruited to bacterial phagosomes via Sulf, that overexpression of Rab7 promotes intracellular growth of strain *Lp02rpsL_WT_* in BMDMs, and that *Lactobacillus pneumophilus* utilizes a lysosomal network for phagosome biogenesis in BMDMs (Li, Fu et al., 2024). Another study revealed that blockade of RAB7 SUMOylation by *S.* Typhimurium ensures the availability of long-lived but functionally compromised RAB7, which is beneficial to the pathogen. These studies demonstrated that SUMO modification of RAB7 exhibits distinct functions for different pathogens (Mohapatra, Gaur et al., 2019). However, RAB-7 was not identified as a SUMOylation substrate in our and other similar proteomics studies in *C. elegans*, which may be attributed to species specificity. An alternative explanation could be that phagosomes that containing bacteria differ from those that containing apoptotic cells, and consequently, RAB7 functions differently in these two processes. We found that, *in C. elegans*, RAB-1 interacts directly with RAB-7, and both RAB-1 knockdown and impaired SUMOylation of GDI-1 led to an increase in the protein content of RAB-7. Aberrant SUMO modification of GDI-1 reduced the interaction between RAB-1 and RAB-7. In addition, when the SUMOylation site of GDI-1 was mutated and PRAF-3 was deleted, the localization of RAB-7 in the endoplasmic reticulum increased, whereas localization in the Golgi apparatus decreased. Normally, RAB-7 can be degraded by lysosomal and proteasomal pathways; however, RAB-7 is degraded only via the proteasomal pathway when the SUMOylation site of GDI-1 is mutated. Based on our findings, we propose that during apoptosis in *C. elegans*, GDI-1 is modified by SUMOylation, which promotes the release of the GDP form of RAB-1. Subsequently, in the presence of PRAF-3, RAB-1-GDP is converted to the GTP form, which then associates with RAB-7, thereby mediating vesicular transport of RAB-7 from the endoplasmic reticulum to the Golgi apparatus to promote the clearance of apoptotic cells (Fig. 8). Our findings demonstrate that SUMO modification of GDI-1 plays a role in apoptotic clearance by regulating RAB-7 through interactions with RAB-1. This study represents the first identification of a function for SUMO modification of GDI-1 in an invertebrate, suggesting the possibility of similar regulatory mechanisms in other organisms.

In mammals, GDI1 is expressed predominantly in neural and sensory tissues, participates in vesicular transport, and may serve as a crucial regulator of several signaling pathways that govern synaptic plasticity, learning acquisition, and memory formation (Lohmer, Clay et al., 2016). Furthermore, a 192P mutation in GDI1 results in X-linked non-syndromic mental retardation in humans (D’Adamo, Menegon et al., 1998). GDI1 expression is elevated in the cerebrospinal fluid of patients with the neurodegenerative disorder Alzheimer’s disease (AD), and inhibition of GDI1 expression attenuates amyloid (Aβ)-induced neurotoxicity in AD (Liu, Liu et al., 2024, More, Kunnecke et al., 2017). We observed that hGDI1 interacted with PIAS1 and GEI-17, and GDI-1 interacted with PIAS1 and GEI-17. Knockdown of pias1 in macrophages resulted in a phagocytosis-deficient phenotype, consistent with that of *C. elegans*. The human genome contains more than 60 Rabs. Aberrant Rab function leads to abnormal distribution of subcellular structures, resulting in neurological disorders, inflammatory diseases, and tumorigenesis. Notably, hGDI1 can interact with RAB1A and RAB-1, and there is also an interactive relationship between GDI-1 and RAB1A and RAB-1. Additionally, the endoplasmic reticulum and Golgi localization of RAB7A in mammalian cells is regulated by RAB1A. Rab1 in mammals has significant functions in signaling, cell migration, and cell surface receptor presentation, and its dysregulated expression is closely associated with human diseases such as cancer, cardiomyopathy, and Parkinson’s disease (Cooper, Gitler et al., 2006, Kiral, Kohrs et al., 2018). Given that hGDI1 interacts with RAB1A in mammals and that RAB7A is also regulated by RAB1A, we determined through sequence comparison analysis that the lysine at position 270 of GDI-1 is relatively conserved in mammalian hGDI1, and the lysine at position 269 of hGDI1 mediated its SUMOylation. The interactions and related functions of these genes are relatively conserved in mammals; therefore, elucidating the regulatory mechanism of SUMO modification of GDI-1 will have significant implications for understanding its role in the pathogenesis of neurodegenerative diseases such as Alzheimer’s disease, in addition to mechanistically revealing its role in apoptotic cell clearance.

In conclusion, we have provided evidence for the regulatory role of GDI-1 SUMOylation in apoptotic cell clearance in *C. elegans* and revealed a previously uncharacterized function of SUMO modification, which expands the known roles of this significant post-translational protein modification.

## Methods

### *C. elegans* strains and genetics

The wild-type strain of *Caenorhabditis elegans* used in this study, as well as the strain used for genetics, was obtained from the Bristol strain N2, which was cultured on plates of Nematode Growth Medium (NGM) containing OP50 at 20 °C. Genetic manipulations were based on Brenner’s research paper “The genetics of *Caenorhabditis elegans*” published in the journal Genetics in 1974 (Brenner, 1974). Nematode strains registered in our laboratory at wormbase were designated SNU (strain) and *xwh* (allele). The mutant alleles used in this study were as follows: linkage group (LG) Ⅰ: *ced-1(e1735)*, *ced-12(n3261)*, *smo-1(ok359)*, and *gei-17(xwh81)*; LG Ⅱ: *mms-21(xwh12)*; LG Ⅲ: *ced-4(n1162)*, and *ced-6(n1813)*; LG Ⅳ: *ced-2(n1994)*, *ced-3(n717)*, *ced-5(n1812)*, *gdi-1(K270R) (xwh89)*, and *praf-3(xwh92)*. The extrachromosomally inherited worm strains obtained by microinjection were subjected to UV irradiation to obtain stable genetically inherited nematode strains. All strains used in this study are listed in the Reagents_Tools_Table.

### Cell lines

Human embryonic kidney 293T (HEK293T) and Mouse L929 cells were obtained from FuHeng Cell Center (Shanghai). Human Jurkat (Clone E6-1), Mouse J774A.1, and RAW264.7 cells were obtained from YiZeFeng Biotechnology (Shanghai) Co., LTD. All cell lines were cultured in DMEM (supplemented with 10% fetal bovine serum and 1% penicillin-streptomycin liquid (100×)) except Jurkat cells, which were cultured in RPMI1640 medium (supplemented with 10% fetal bovine serum and 1% penicillin-streptomycin liquid (100×)). The cells were then incubated at 37 °C with 5% CO_2_. All cell lines used in this study are listed in the Reagents_Tools_Table.

### Mouse

Animal protocols were approved by Academic Committee of Shaanxi Normal University. All mice were cared for according to the Shaanxi Normal University guidelines for the care and use of laboratory animals, and were in good general health based on appearance and activity. The strains of mice used in the experiments were Balb/c and C57BL/6J. The experimental mouse strains were kept in the Animal Experiment Center of Shaanxi Normal University according to standard methods.

### RNAi experiments

RNA interference (RNAi) refers to the introduction of double-stranded RNA (dsRNA) into hermaphroditic *C. elegans*, which leads to sequence-specific and rapid degradation of endogenous mRNAs in order to knock down gene expression. The expression of target genes can be effectively blocked by RNAi and gene silencing can be transmitted between tissues throughout the adult and its progeny. We used a food-fed RNAi approach (Chen et al., 2010), which was performed by feeding *E. coli* HT115 with the corresponding RNAi vector for worm target genes. For genetic analysis, 10-20 adult worms were placed on NGM plates containing 1 mM isopropyl β-d-thiogalactopyranoside (IPTG) and 100 µg/mL ampicillin containing RNAi bacteria, and the apoptotic phenotypes of their progeny were counted. For biochemical experiments, 50-100 worms at different periods were placed on NGM plates containing RNAi bacteria, and the worms were collected with M9 buffer after 48 h. Proteins were extracted for subsequent experiments.

### Quantification of Embryonic and Germline Cell Corpses

A control group was required for each RNAi treatment to confirm the achievement of RNAi effect and the confidence of the data. The number of “button-like” shaped apoptotic cells in the head region of living embryos was counted using Nomarski DIC optics at different developmental stages (comma, 1.5 Fold, 2 Fold, 2.5 Fold, 3 Fold, and 4 Fold), with 15 embryos per nematode strain counted at each developmental stage. To determine the number of apoptotic germ cells, apoptotic cells were counted in the meiotic region of one gonadal arm of at least 15 nematodes at 12, 24, 36, 48, and 60 h after the L4 stage. The average number of embryonic and germline cell corpses was compared with those of other transgenic worms using an unpaired *t*-test.

### CRISPR/Cas9-mediated genome editing

The Cas9 editing involved in this project is a more mature method studied by previous researchers, in which the worm genome is edited to obtain specific allele knockout mutants, in situ insertions, or fixed-point mutations in *C. elegans* (Paix, Wang et al., 2014). Briefly, target gene sgRNAs were analyzed at http://crispr.mit.edu. 1-3 sgRNAs were selected to ensure the knockdown or knock-in efficiency. The sgRNA sequences were introduced by PCR into the Cas9-expressing vector *pDD162-peft-3-cas9-sgRNA* empty. *dpy-10* sgRNA plasmid 20 ng/μl, target sgRNA plasmid 20 ng/μl, target ssODN 2μM, *dpy-10* ssODN 2μM co-injected into 50 worms. The progeny were then identified by single-worm PCR to identify which were successfully edited. The target sgRNA and the repairing oligo sequences used in this study are listed in the Reagents_Tools_Table

### Time-lapse Imaging and Microscopy

The method was referred to the previous article with modifications (Gan, Wang et al., 2019). Worms were placed in M9 buffer containing 2 mM levamisole, covered with a coverslip, and sealed with beeswax to keep them in liquid. Thirty germ cell corpses were tracked and photographed in each experiment. Germ cell corpses were tracked using a ZEISS Imager M2 orthogonal fluorescence microscope and recordings were taken every 5-10 min. During the experiment, worms were constantly checked for survival to ensure data reliability. To record the duration of embryonic cell corpses, early embryos (2-cell stage) were placed in egg salt buffer (118 mM NaCl and 48 mM KCl) and mounted on slides with agar pads. Images in a 25-section Z-series (1 μm/section) were captured every 1 min for 400 minutes using a ZEISS Imager M2 orthogonal fluorescence microscope.

Worms tagged with fluorescence were imaged by DIC and fluorescence using an Axio Imager M2 microscope (ZEISS). Images were processed and viewed using ZEN 2 pro software (ZEISS). Immersol 518F oil (Zeiss) was used. All images were captured at 20 °C.

To monitor the uptake and degradation of cell corpse chromatins in phagosomes and secreted soluble (ss)GFP in coelomocytes, worms were observed using fluorescence microscopy at the indicated times. Secreted soluble (ss)GFP worms (*arIs36*) were heat-shocked for 1 h at 33 °C and continued to grow at 20 °C.

### Quantification of Phagosomal Markers

Transgenic worms with fluorescent markers genes for each stage of phagosome (CED-1::GFP, GFP::RAB-5, GFP::RAB-7, mCHERRY::RAB-7, LAAT-1::GFP, LAAT-1::mCHERRY, NUC-1::mCHERRY and CPL-1::mChiOnt) were utilized for the relevant indicated strains, and clinical slides were prepared by picking the treated worms on slides with agar. One hundred apoptotic cells were counted for each worm species, and the apoptotic cells were observed under the fluorescence of ZEISS Imager M2 microscope to determine whether the apoptotic cells were surrounded by fluorescent phagosome marker genes. The fluorescent-labeling ratio of phagosomal genes is the number of fluorescently labeled apoptotic cells divided by the total number of apoptotic cells.

### Acridine Orange (AO) Staining

To detect phagosomal acidification by AO, adult worms were incubated in M9 medium containing 0.1 mg/ml AO and a small amount of OP50 in the dark for 1 h at room temperature. After treatment, worms were transferred to NGM plates at room temperature in the dark for 2 h and mounted on 2% agar pads. Images were captured using DIC and fluorescence microscopy. At least one hundred apoptotic cells were counted in each strain.

### Immunoprecipitation and LC-MS/MS analysis

The collected unsynchronized wild-type and transgenic worms (N2, *ha::smo-1*, and *ha::smo-1; P_ced-1_gdi-1::gfp*) were resuspended with lysis buffer (25 mM Tris-HCl, pH 7.4, 150 mM NaCl, 0.1% sodium deoxycholate, 1% NP-40, 10% glycerol, 1× mixture of protease inhibitors [Complete, EDTA free], and 1 mM PMSF), homogenized completely with tissue grinders, and centrifuged at 20,000 × *g* at 4 °C for 30 min. 4 mg proteins were co-incubated with the Anti-HA Magnetic Beads (Thermo Scientific, Cat#88837) for 6 h at 4 °C under rotary mixing. The bound proteins beads were rinsed four times with TBS wash buffer (50 mM Tris-HCl, pH 7.6, 300 mM NaCl). The beads were then resuspended in 50 µl 2 × SDS loading buffer and boiled for 5 min at 100 °C for western blotting.

After immunoprecipitation, the samples from three replicate experiments were mixed together and identified by mass spectrometry (HOOGEN Biological Co. Shanghai). Mass spectrometry was performed as previously described (Yuan et al., 2022a).

### Western Blot Analysis

HEK293T cells and worms were resuspended in lysis buffer containing a protease inhibitor cocktail and homogenized completely using a sonic crusher or tissue grinder. Homogenized samples centrifuged at 20,000 × *g* for 15 min at 4 °C to remove debris and protein concentration was quantified by BCA method, then 30 µg samples were boiled for 5 min at 100 °C in 5× SDS loading buffer for further analyses. Proteins were loaded onto SDS-PAGE, transferred onto PVDF membranes, blocked with 5% skimmed milk, and incubated overnight with the primary antibodies (β-actin loaded control) at 4 °C. After washing with TBS buffer, probed with HRP-conjugated secondary antibodies, and developed with the Immobilon^®^ Western chemiluminescent HRP substrate (Millipore). Protein bands were visualized using a MiniChemi610 (SAGE creation). All antibodies used in this study are listed in the Reagents_Tools_Table.

### Yeast two-hybrid analysis

Genes indicated for detecting protein interactions were cloned into the pGADT7 and pGBKT7 vectors. Y2HGold was cultured in liquid YPDA medium supplemented with 0.25 μg/ml kanamycin and 40% glucose, when the OD_600_ was shaken to 0.3-0.6, prepared for the Y2HGold chemically competent cell. 240 μL 50% PEG3350, 36 μL 1 M LiAC, 10 μL Salmon Testes DNA, and 0.7 μg pGADT7 and pGBKT7 plasmid were added sequentially into Y2HGold chemically competent cell. Transformed Y2HGold chemically competent cell were coated on SD-Leu-Trp medium for 3-5 days at 30 °C. Single colonies were transferred to SD-Leu-Trp liquid medium at 280 rpm overnight in a shaker at 30 °C, and the yeast solutions were dropped onto SD-Leu-Trp and SD-Leu-Trp-His plates to test for interactions.

### Co-IP assay

Co-immunoprecipitation experiments were performed as previously described (Yuan et al., 2022a), with some modifications. To detect protein interactions, HEK 293T cells in a 10 cm plate were transfected with 8 µg of each plasmid and 48 µl 1µg/µl PEI (Polysciences, Inc., 24765-1). 48 hours after transfection, the cells were lysed with cell lysis buffer (25 mM Tris-HCl, pH 7.6, 150 mM NaCl, 10mM MgCl_2_, 1% NP-40, and EDTAfree protease inhibitors), sonicated for 15 s, and centrifuged at 20,000 × *g* at 4 °C for 20 min to remove debris. 2 mg total protein of was incubated with anti-HA (Thermo, 88837) or anti-C-MYC (Thermo, 88842) magnetic beads at 4 °C for 2-3 h. The precipitates were 1×SDS loading buffer with 5% 2-mercaptoethanol boiled for 5 min at 100 °C. The samples were analyzed by WB.

To detect the interactions of GDI-1 with RAB-1 and RAB-1 with RAB-7, unsynchronized worms raised on 10 NGM plates were harvested. The samples were resuspended in lysis buffer containing a protease inhibitor cocktail, homogenized completely with tissue grinders, and centrifuged at 20,000 × *g* at 4 °C for 30 min to remove debris for Co-IP. Supernatants were incubated with anti-FLAG M2 affinity gel (Sigma-Aldrich, A2220) or Protein A/G Mix Magnetic Beads (Millipore, LSKMAGAG) at 4 °C for 4-6 h. When used with Protein A/G Mix Magnetic Beads, lysates were first incubated with primary antibody (GDI-1 or RAB-1 polyclonal antibody) at 4 °C for 2-4 h. The precipitates were boiled in 1×SDS loading buffer with 5% 2-mercaptoethanol and analyzed using WB.

### Purification of Recombinant Proteins and GST Pull Down

Plasmids expressing each protein were generated using GST or HIS tags. The recombinant proteins GST, GST::ARL-8, GST::GDI-1, GST::GEI-17, GST::GDI-1(K269R), GST::RAB-1, GST::RAB-7, and GST::hGDI1 were expressed in *E. coli* BL21 strain induced by IPTG. GST-tagged proteins in BL21 were lysed in GST lysis buffer (25 mM Tris-HCl, pH 7.6, 150 mM NaCl, 0.5% NP-40) with 1 mM PMSF and 1 mg/ml lysozyme and sonicated until the lysates were clear. GST-tagged proteins were incubated with Glutathione Sepharose 4 B (GE Healthcare, 17-0756) beads for 2-3 h at 4 °C, washed three times with GST lysis buffer, and eluted with 20 mM or 30 mM glutathione reduced (AMERSCO, 0399). 6HIS::MYC::MMS-21::6HIS, 6HIS::MYC::ARL-8::6HIS, 6HIS::MYC::GDI-1::6HIS, 6HIS::MYC::GDI-1(1-150 aa)::6HIS, 6HIS::MYC::GDI-1(150-444 aa)::6HIS, 6HIS::MYC::RAB-1::6HIS, 6HIS::MYC:RRAB-28::6HIS, 6HIS::MYC::RABY1::6HIS, 6HIS::HA::RAB-7::6HIS, 6HIS::HA::PRAF-3::6HIS, 6HIS::MYC::PIAS1::6HIS, and 6HIS::MYC::RAB1A::6HIS were expressed in *E. coli* BL21 (DE3) strain induced by IPTG. HIS-tagged proteins were lysed using the same approach for the GST-tagged proteins. After sonication, the HIS-tagged proteins were incubated with Ni-NTA Superflow (QIAGEN) beads for 2-3 h at 4 °C and eluted with 10/30/100/150 mM imidazole (Merck) in GST lysis buffer. Purified proteins were stored in GST lysis buffer containing 20% glycerol at -80 °C.

GST pull-down experiments were performed as previously described (Yuan et al., 2022a) with some modifications. 2 µg purified GST-tagged proteins immobilized on Glutathione Sepharose 4 B beads were incubated with 0.5 µg HIS-tagged proteins in binding buffer (25 mM Tris-HCl, pH 7.4, 150 mM NaCl, 0.1% NP-40) for 2 h at 4 °C. The resins were washed three times in washing buffer (25 mM Tris-HCl, pH 7.4, 300 mM NaCl, 0.1% NP-40), and the bound proteins were analyzed by WB.

### SUMOylation of GDI-1 *in vitro* and *in vivo*

SUMOylation experiments *in vitro* were performed as previously described (Kaminsky, Denison et al., 2009), with some modifications. To examine SUMOylation *in vitro*, 2 µg HA::GDI-1 or HA::GDI-1(K270R), 0.2 µg hSAE1/SAE2, 0.5 µg hUBC9, and 4 µg MYC::hSUMO1 were incubated in SUMOylation buffer (20 mM HEPES, pH 7.4, 100 mM NaCl, 5 mM MgCl_2_, 2 mM ATP) at 37°C at the indicated time points to allow formation of soluble oligomers. The reactions with the control and HA::GDI-1(K270R) were incubated for 1 h at 37 °C. To confirm the SUMOylation of GDI-1 *in vivo*, unsynchronized worms (*ha::smo-1*, *flag::gdi-1*, *ha::smo-1; flag::gdi-1*, *flag::gdi-1(K270R)*, and *ha::smo-1; flag::gdi-1(K270R)*) were collected and lysed in lysis buffer (25 mM Tris-HCl, pH 7.4, 150 mM NaCl, 0.1% sodium deoxycholate, 1% NP-40, 10% glycerol) containing 1× mixture of protease inhibitors [Complete, EDTA free] and 20 mM *N*-Ethylmaleimide (Sigma-Aldrich, E3876). Protein (4 mg) was immobilized on an anti-FLAG M2 affinity gel at 4 °C for 4 h, washed three times with washing buffer (25 mM Tris-HCl, pH 7.6, 300 mM NaCl) and analyzed by WB.

### Lifespan Assay

Lifespan experiments were performed as previously described (Wan, Yuan et al., 2021), with some modifications. Lifespan assays by *P. aeruginosa* PA14 infection were conducted at 25 °C, and *S. typhimurium* SL1344 infection were conducted at 20 °C. The L4 stage was designated as day 0, at least 100 worms were used for each experiment, and animals were transferred every 1-2 days to fresh plates to eliminate overcrowding by progeny until they laid no further eggs. Worms were marked as dead when they did not move, pump, or respond to the stimuli. Survival curves were plotted using GraphPad Prism 8.

### Yeast three-hybrid assay

The method was referred to the previous article with modifications (Gao et al., 2019). The yeast three-hybrid assay in this subject was used to detect GDI-1 and PRAF-3 competitive binding to RAB-1. *gdi-1* alone, or together with *praf-3*, were cloned into pBRIDGE. *rab-1* was cloned into pGADT7. The pGADT7 and pBRIDGE plasmids were transformed into the Y2HGold strain and cultured on SD-Leu-Trp double-dropout solid culture medium. On the SD-Leu-Trp-His dropout medium, GDI-1 interacted with RAB-1 on SD-Met-Leu-Trp-His dropout medium to induce the expression of PRAF-3 and test for competitive binding.

### Extraction of membrane proteins

Membrane proteins extraction was performed using a Membrane Protein Extraction Kit (Bestbio, BB-3103). Unsynchronized N2, *gdi-1(K270R)*, and *praf-3* worms were collected and lysed with 400 μL Reagent A from the kit, and incubated at 4 °C for 25 min with rotation. After centrifugation, the supernatant was incubated at 37 °C for 8 min and centrifuged again, after which the supernatants containing cytoplasmic proteins and the lower layers containing membrane proteins were collected. And the membrane proteins were resuspended by adding Reagent B. An ice bath was performed for 2 min, followed by a water bath at 37 °C for 5 min. After centrifugation, 50 μL Reagent C was added to dissolve the lower layer of solution, which is the membrane proteins. Proteins were quantified using the bicinchoninic acid method and analyzed by WB.

### Extraction of Endoplasmic Reticulum and Golgi proteins

Extraction of ER and Golgi proteins were performed using Endoplasmic Reticulum Protein Extraction Kit (Solarbio, EX1260) and Golgi Protein Extraction Kit (Solarbio, EX1240). Unsynchronized N2, *gdi-1(K270R)*, and *praf-3* worms or control/*rab1a* knockdown HEK 293T cells were washed with cold PBS. For the extraction of ER, samples dissolved in 400 μL of Reagent A were homogenized using a homogenizer or sonication. After centrifugation at 1,000 × *g* for 5 min at 4 °C, the supernatants were centrifuged again at 11, 000 × *g* for 10 min. Subsequently, centrifugation at 50,000 × *g* for 45 min was performed, and the precipitates were resuspended in Reagent B and centrifuged again at 50,000 × *g* for 45 min. Finally, the precipitates were resuspended in 50 μL of protein extract C, which is an endoplasmic reticulum protein.

For the extraction of Golgi, samples dissolved in 400 μL of Reagent A were homogenized using a homogenizer or sonication. The samples were centrifuged at 3,000 × *g* for 10 min and 10 μL of Reagent WT was added. The samples were centrifuged at 20,000 × *g* for 20 min, and 500 μL of Reagent B was added to the precipitates and mixed thoroughly. After centrifugation at 20,000 × *g* for 30 min, 50 μL of Reagent C were added to the precipitates, which were Golgi proteins. The extracted endoplasmic reticulum and Golgi proteins were quantified by BCA method and analyzed by WB.

### Antibodies

GDI-1, RAB-1, SCA-1, TBB-1, RAB-7, CNX-1, and SYX-6 antibodies were generated in mice by injecting recombinant proteins. BALB/c mice at 8-12 weeks were intraperitoneally injected with 80 μg of purified protein in a 1:1 mixture with Freund’s Adjuvant, Complete (Sigma-Aldrich, F5881). After two weeks, mixtures of 80 μg of purified protein with Freund’s Adjuvant, Incomplete (Sigma-Aldrich, F5506) were intraperitoneally injected at two-week intervals for a total of three injections. Mice were blooded from eye sockets and sera were collected for antibody detection.

### Quantitative Real Time PCR

Quantitative real-time PCR experiments were performed as previously described (Yuan et al., 2022a) with some modifications. RNA was extracted from the cells by chloroform extraction, followed by ethanol precipitation and DNase treatment. Complementary DNA was synthesized from 1 μg of RNA using HiScript® III RT SuperMix for qPCR (+ gDNA wiper) (Vazyme, CAT# R323) according to the manufacturer’s instructions. Quantitative RT-PCR was performed using the ChamQTM Universal SYBR® qPCR Master Mix (Vazyme, CAT# Q711).

### shRNA-mediated Gene Silencing and Lentivirus Packaging

The targeted shRNAs were cloned into pGREEN-puro plasmids. Scrambled RNA control and shRNAs were transfected into cells using electrotransfer or lentivirus. After 48 h, the cells were used for subsequent analyses.

For lentiviral packaging, 293 TN cells were co-transfected with the targeted shRNA pGREEN-puro plasmid and pSBH155/156 packaging plasmids using PEI. After 24 h, the media were changed to DMEM (supplemented with 10% FBS and 1% penicillin-streptomycin liquid (100×)), and the cell culture fluids were harvested at 48h and 72h after infection, respectively which were the lentivirus infection solutions.

### Bone Marrow-Derived Macrophages

Bone marrow cells were isolated from 8 weeks C57BL/6J mice. BMDM were differentiated and cultured for one week in DMEM supplemented with 10% fetal bovine serum, 1% penicillin, and 20% L929 cell supernatant. The cells were then incubated at 37 °C with 5% CO_2_.

### Electrotransfer

Mouse RAW264.7 cells transfection were performed using Neon^TM^ Transfection Kit (Invitrogen, MPK10096) by Invitrogen Neon^TM^ Transfection System. RAW264.7 cells were rinsed with PBS and resuspended with R Buffer, 100 μL 4×10^6^ RAW264.7 cells and 6 μg plasmids were transfected under electric shock conditions at 1700 V, 25ms, 1 pules. Transfected cells were cultured in DMEM containing 10% FBS, and after 48 h of incubation, subsequent experiments were performed.

### Efferocytosis assay and Flow Cytometry

Jurkat cells were stained with 1 μM CellTracker™ Red CMTPX (Invitrogen, E34552) for 30 min, and then irradiated with 150 J UV, followed by incubation at 37 °C 5% CO_2_ for 3 h to generate apoptosis. The apoptotic Jurkat cells were resuspended in 1x PBS, and added to RAW264.7, J774A.1, or BMDM cells transfected with the *pias1-shRNA* at a ratio of 10:1, followed by incubation at 37 °C for 6 h. After incubation, the macrophages were rinsed twice with PBS to remove unbound apoptotic Jurkat cells.

For microscopic imaging, 1 ml of DMEM were added to the cell culture dish. Apoptotic Jurkat cell uptake was quantified as the percentage of Apoptotic Jurkat cell-positive macrophages out of the total number of macrophages per field of view. For flow cytometric experiments, the macrophages were digested with trypsin, resuspended in 200 µL PBS, and analyzed on a BECKMAN COULTER CytoFLEX S flow cytometer. Data analysis was performed using CytExpert software.

### Statistical analyses

Statistical analyses were performed using Microsoft Office Excel, and bar graphs or other statistical plots were produced using GraphPad Prism Software. Significant differences between samples appearing in the paper were tested using a two-tailed unpaired *t*-test, with error bars denoting SEM. *, p < 0.05; **, p < 0.01; ***, p < 0.001.

## Acknowledgments

We thank Drs. Chonglin Yang and the *C. elegans* Genetic Center for *C. elegans* strains, and Dr. Cheng-Gang Zou for providing *P. aeruginosa* PA14 and *S. typhimurium* SL1344 strains. This work was partially supported by the National Natural Science Foundation of China (Grant No. 31871387 to Hui Xiao), National Natural Science Foundation of China (Grant No.32370799 to Hui Xiao), National Natural Science Foundation of China Youth Program (Grant No.32300622 to Lei Yuan), Sanqin Bochuang Talent Support Program of Shaanxi Province (Grant No. 2024SQBC004 to Lei Yuan), Natural Science Foundation of Shaanxi Province (Grant No.2024JC-YBMS-638 to Hui Wang), the program of Innovative Research Team for the Central Universities (Grant No. GK202302003 to Hui Xiao).

## Author contributions

**Lei Yuan**: Conceptualization; Data curation; Formal analysis; Funding acquisition; Investigation; Writing—original draft. **Peiyao Li**: Conceptualization; Data curation; Formal analysis; Investigation; Writing—original draft. **Aiying Ma**: Conceptualization; Data curation. **Chao Li**: Data curation. Fuqiao Liu: Data curation. **Fuqiao Liu**: Data curation. **Yunmin Xie**: Data curation. **Qian Zheng**: Data curation. **Hui Wang**: Conceptualization; Supervision; Funding acquisition; Writing—original draft. **Hui Xiao**: Conceptualization; Supervision; Funding acquisition; Methodology; Writing— original draft.

---

**Figure EV1.**
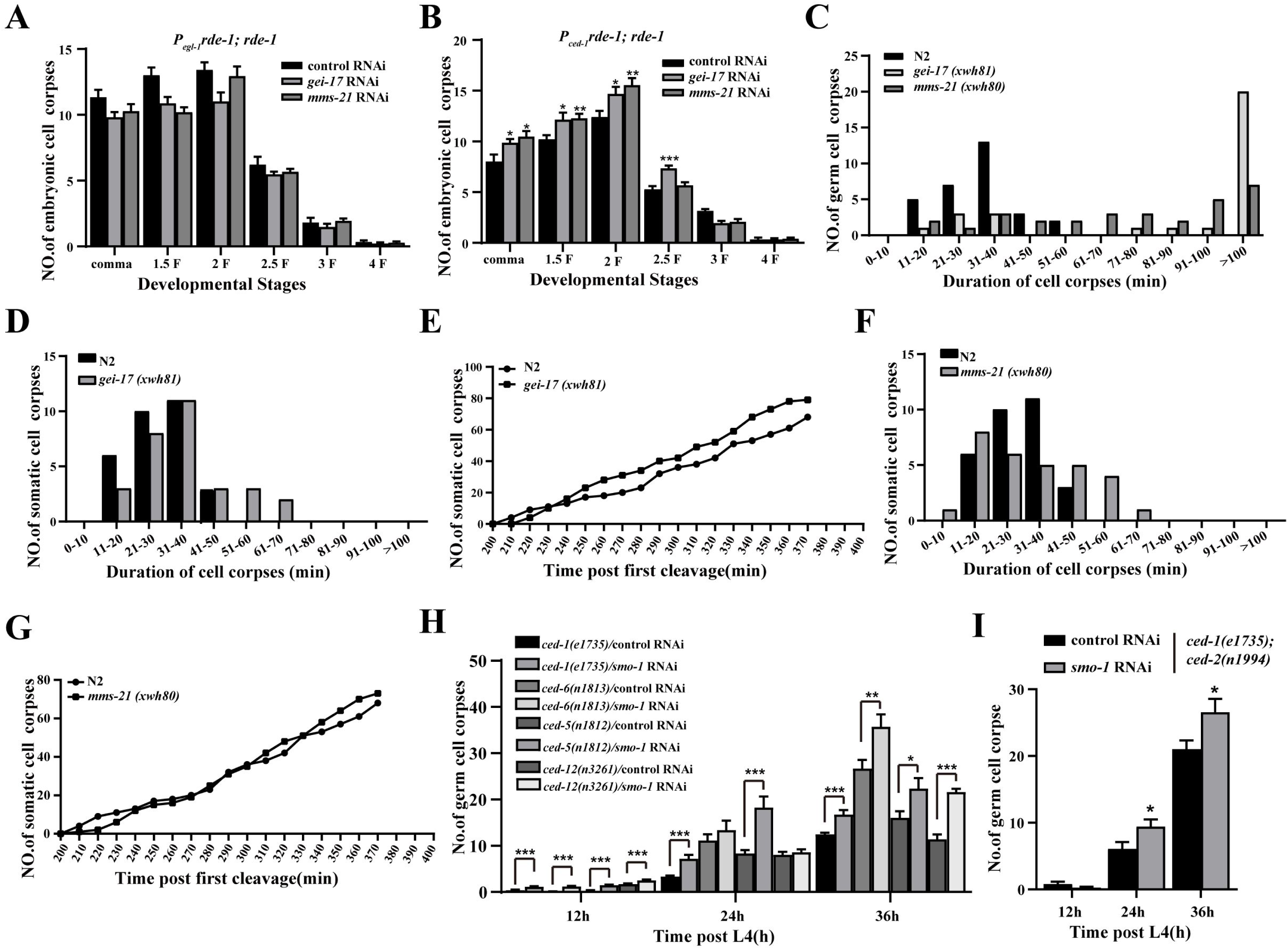
SUMO modifications functions independently of either pathway for cell corpse removal. (A and B) Quantification (mean ± SEM) of embryonic cell corpses at different embryonic stages using the indicated strains. Fifteen embryos were scored for each embryonic stage for each strain. (C) Four-dimensional microscopy analysis of germ cell corpse duration was performed in N2, *mms-21 (xwh80)*, and *gei-17 (xwh81)*. The persistence of 30 germ cell corpses was also monitored. (D and F) The number of embryo corpses was measured by four-dimensional microscopy in N2, *gei-17 (xwh81)*, (D), and *mms-21 (xwh80)* (F). Embryos (2-cell stage) were isolated from adult worms and imaged every minute for 400 min. (E and G) Four-dimensional microscopy analysis of somatic cell corpse persistence in N2, *gei-17 (xwh81)*, (E), and *mms-21 (xwh80)* (G). Thirty somatic cell corpses were analyzed for each genotype. (H and I) Quantification (mean ± SEM) of germ cell corpses from the indicated strains. Cell corpses of each animal were scored in fifteen animals at every time point (h post L4) as indicated. The unpaired *t* test was performed in this figure. *P < 0.05, **P < 0.01, ***P < 0.001. All error bars indicate the mean ± SEM.

**Figure EV2.**
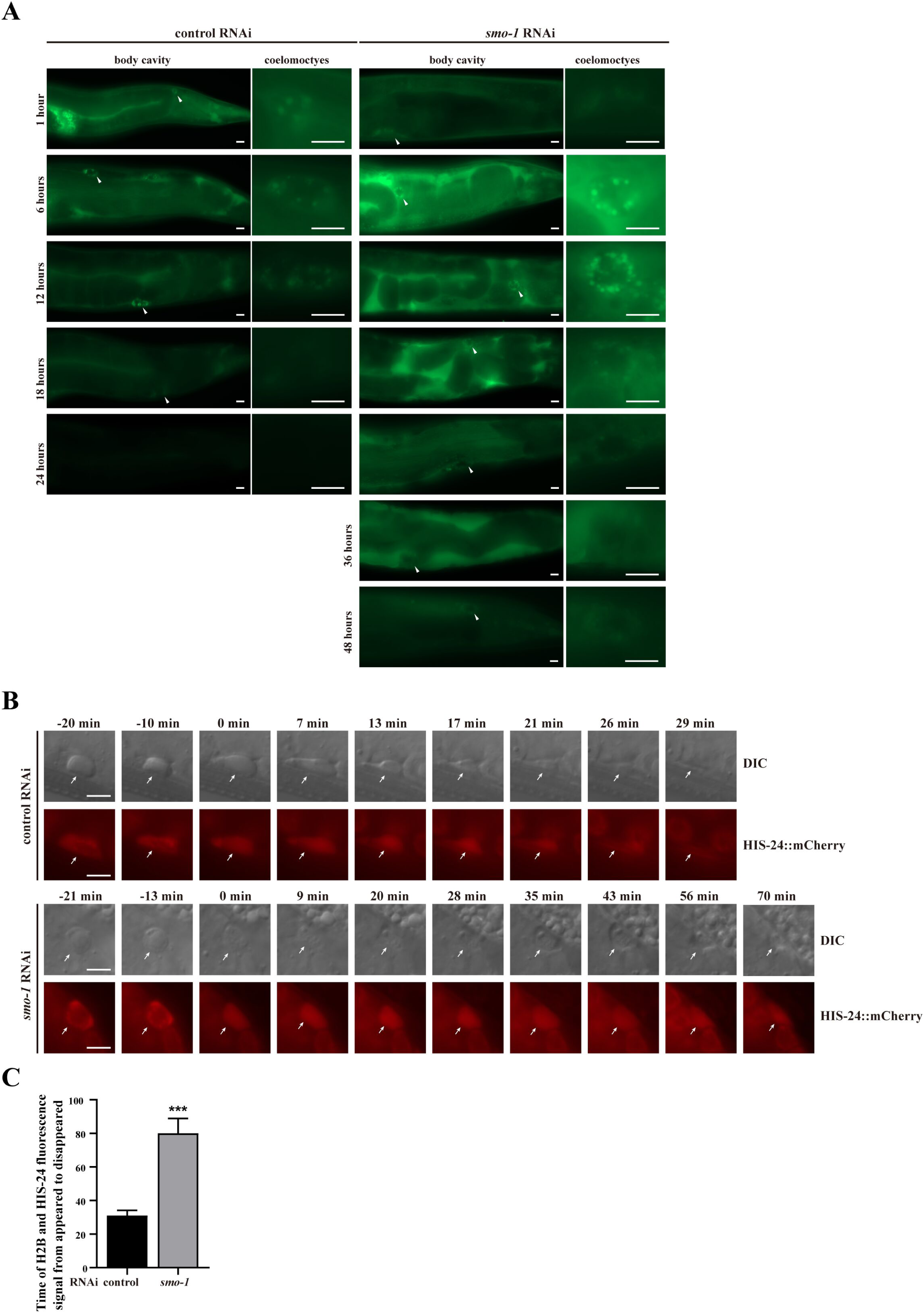
SUMOylation regulates phagosome degradation. (A) Animals expressing ssGFP controlled by a heat-shock promoter treated with control and *smo-1* RNAi were heat shocked for 60 min at 33°C, and the uptake and degradation of ssGFP in coelomoctyes were monitored at the indicated time points. Left, accumulation of ssGFP in the body cavity (400×), and arrows indicate coelomocytes that are enlarged in the right pictures (1000×). Bars, 10 μm. (B) Time-lapse chasing of HIS-24::mCherry positive phagolysosomes in N2/control and *smo-1* RNAi worm germlines. The time point at which the HIS-24::mCherry mostly rings the cell corpse was set to 0 min. Arrows indicate the continuous presence of cell corpses (DIC, top row) and HIS-24::mCherry (fluorescence, bottom row). DIC and fluorescence images were obtained at certain time intervals. Bars, 5 µm. (C) The persistence of the HIS-24::mCherry fluorescence signal in five germ cell corpses in N2/control and *smo-1* RNAi worms was monitored. An unpaired *t* test was performed in this figure. ***P < 0.001. All error bars indicate the mean ± SEM.

**Figure EV3.**
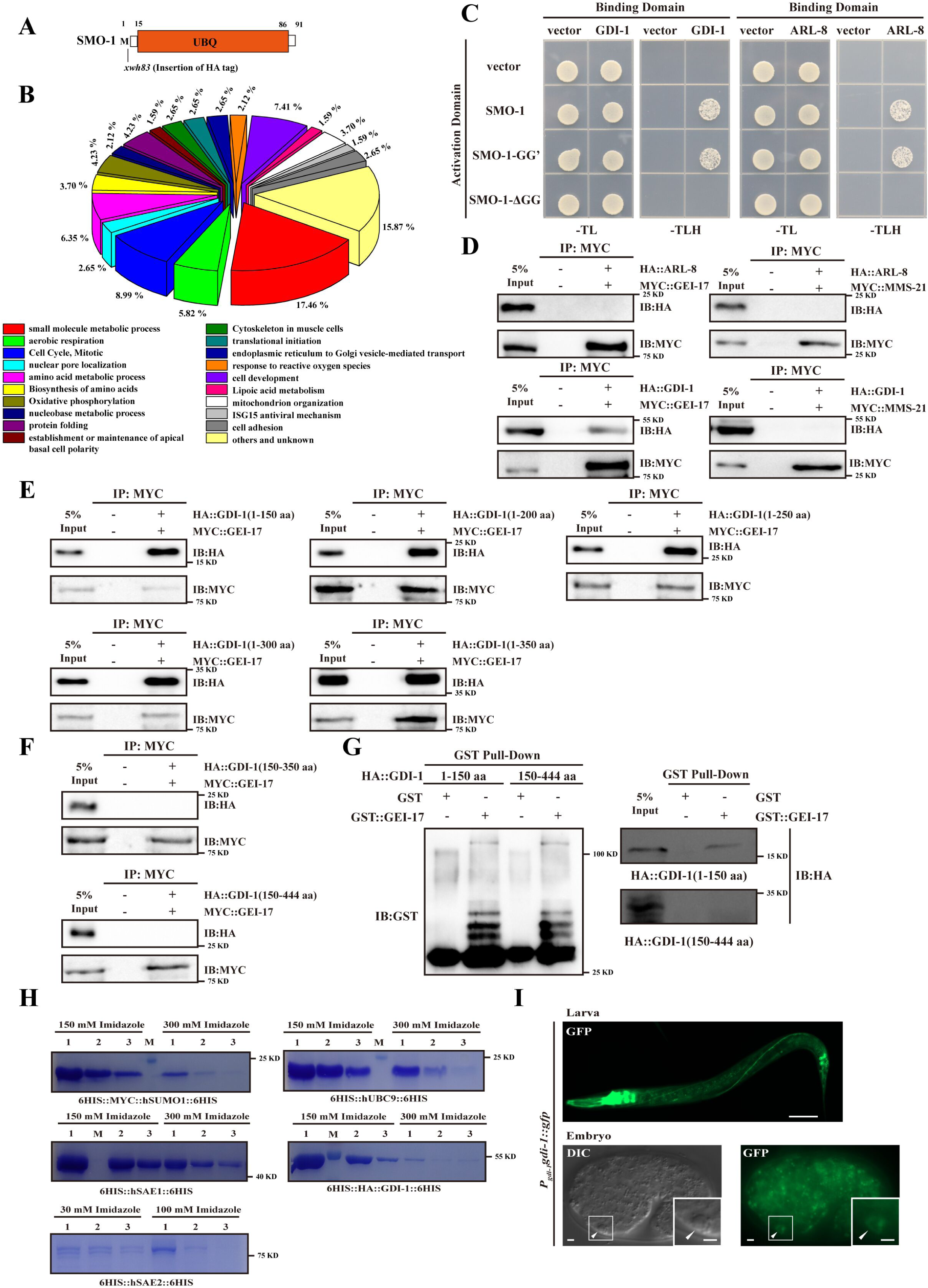
Identification of GDI-1 as a potential substrate of GEI-17 for apoptotic cell clearance. (A) Schematic illustration of the HA tag insertion generated by CRISPR-Cas9 editing of endogenous *smo-1* loci. Amino acids near the insertion site are indicated. (B) Grouping of identified SUMO targets into broad functional classes. (C) Yeast two-hybrid assay of the interactions between GDI-1 or ARL-8 and SMO-1 WT/GG’/ΔGG. SMO-1 GG’, residues after C-terminal di-glycine were deleted. SMO-1 ΔGG, di-glycine residues in SMO-1 were deleted. -TL, medium lacking Trp and Leu; -TLH, medium lacking Trp, Leu, and His. (D) The interactions between ARL-8 or GDI-1 and GEI-17 or MMS-21 were examined by CO-IP in 293T cells. (E) Coomassie brilliant blue (CBB) stained gel of MYC::hSUMO1, hSAE1, hSAE2, hUBC9, and HA::GDI-1 following affinity purification using Ni-NTA agarose. (F) GDI-1::GFP localization in larvae (top) and embryos (bottom) of xwhIs (*Pgdi-1gdi-1::gfp*) worms were imaged by the gfp channel using an Imager M2 microscope. Arrows indicate cell corpses in embryo images. The boxed region is magnified (2.0×) in the inset. Scale bars: 50 µm (top) and 2 µm (bottom). (G-I) The interaction between truncation of the GDI-1 amino acids and GEI-17 was examined by CO-IP in 293T cells (G and H) and GST pulldown assays (I).

**Figure EV4.**
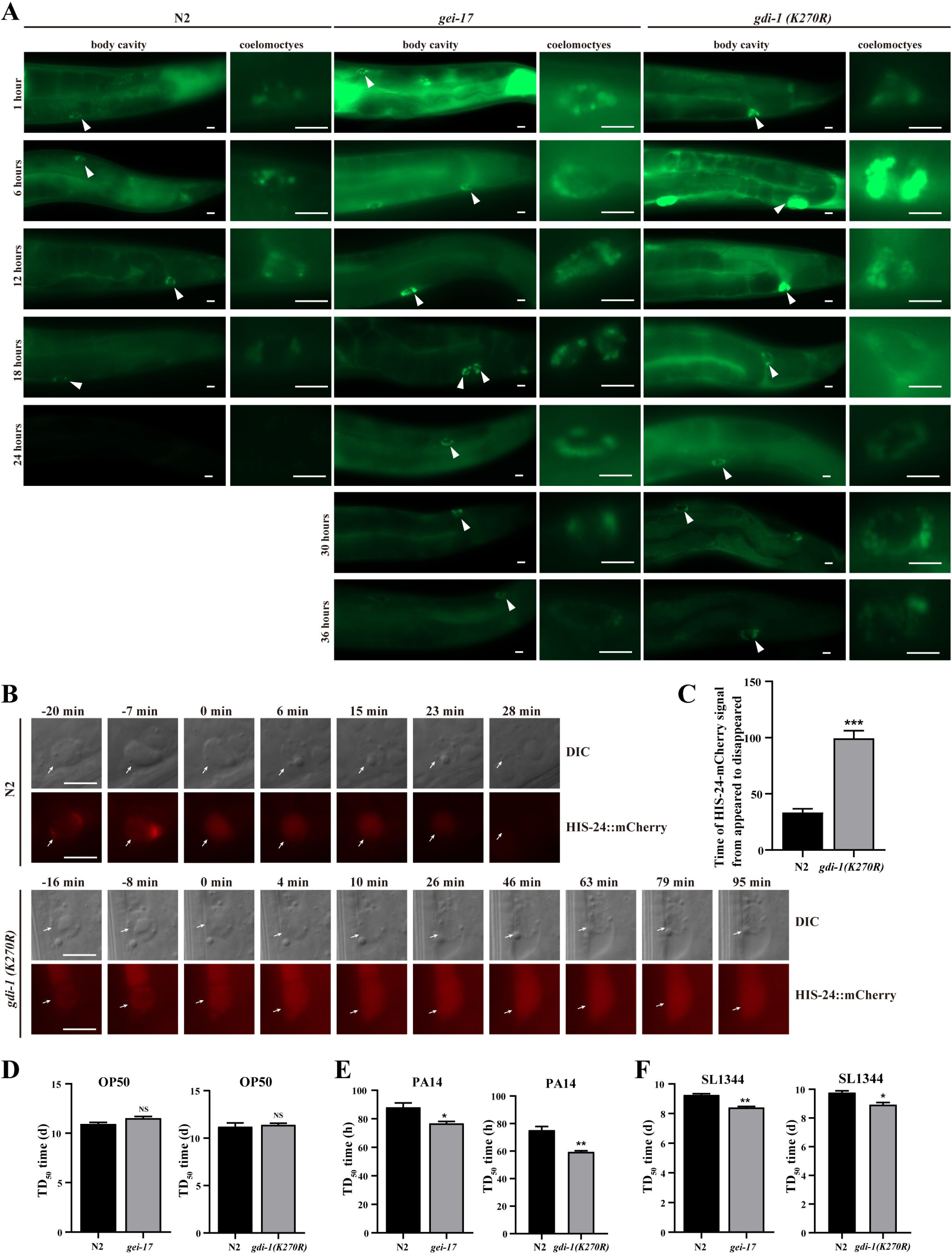
GDI-1 SUMOylation at K270 affects Endosomal/Lysosomal degradation and phagosome degradation. (A) N2, *gei-17*, and *gdi-1(K270R)* animals expressing ssGFP controlled by a heat-shock promoter were heat shocked for 60 min at 33°C, and the uptake and degradation of ssGFP in coelomoctyes were monitored at the indicated time points. Left, accumulation of ssGFP in the body cavity (400×), and arrows indicate coelomocytes that are enlarged in the right pictures (1000×). Bars, 10 μm. (B) Time-lapse chasing of cell corpses in DIC and HIS-24::mCherry positive phagolysosomes in N2 and *gdi-1(K270R)* germlines. The time point at which the HIS-24::mCherry ring was first detected on the cell corpses was set as 0 min. Bars, 5 µm. (C) The persistence of the HIS-24::mCherry fluorescence signal in five germ cell corpses in N2 and *gdi-1(K270R)* worms was monitored. (D-F) Time to 50% mortality of *C. elegans* in OP50 (D), PA14 (E), and SL1344 (F). An unpaired *t* test was performed in this figure. *P < 0.05, **P < 0.01, ***P < 0.001. All error bars indicate the mean ± SEM.

**Figure EV5.**
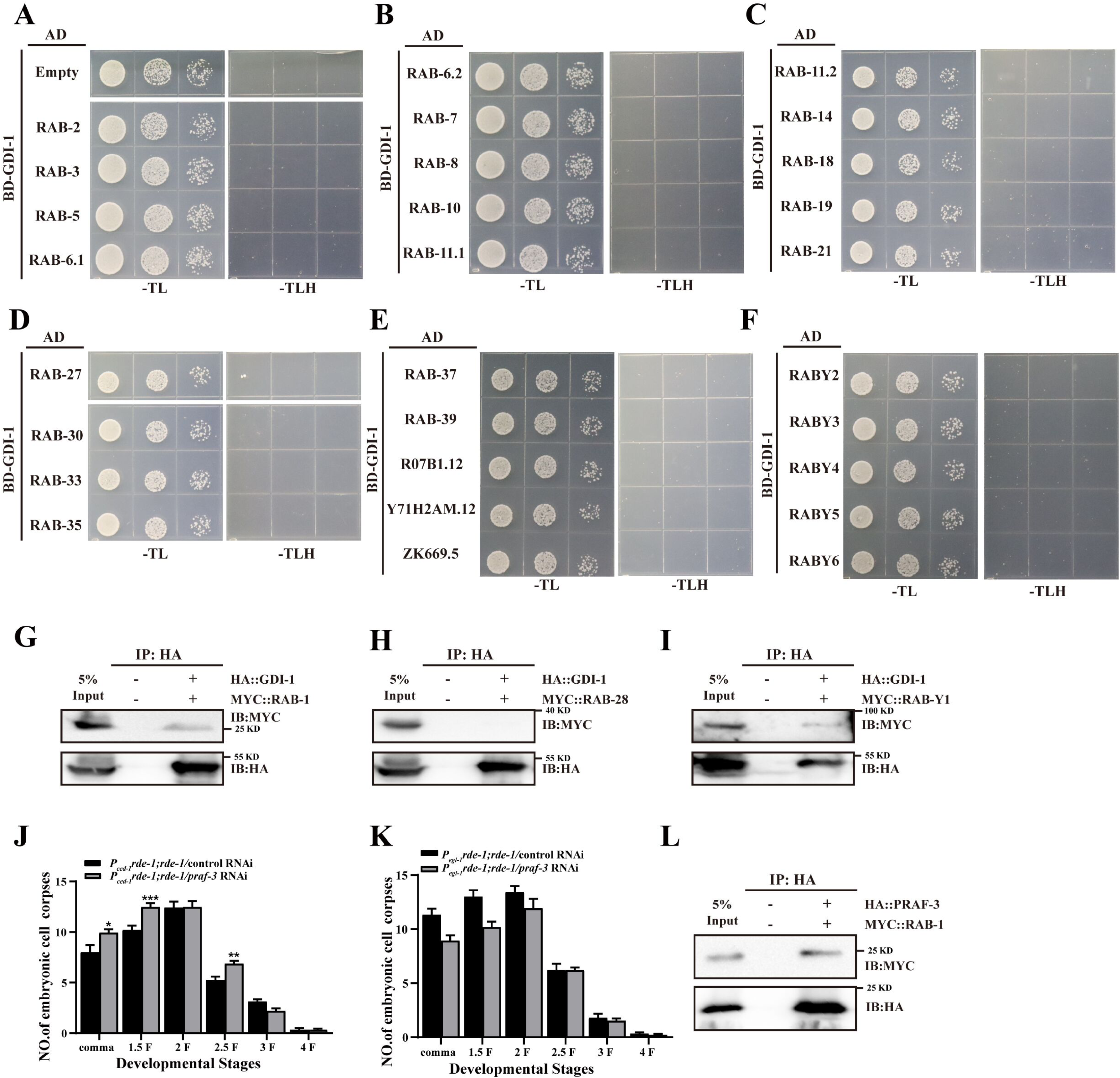
Identification of RAB-1 as an interaction protein for GDI-1. (A-F) Yeast two-hybrid assay for screening RABs that interact with GDI-1. (G-I and L) The interaction between GDI-1 and RAB-1 (G), GDI-1 and RAB-28 (H), GDI-1 and RAB-Y1 (I), and PRAF-3 and RAB-1 (L) were examined by CO-IP in 293T cells. (J and K) Different stages (h post L4) of germ cell corpses were quantified (mean ± SEM) in the indicated strains. Fifteen adult worms were scored at each stage for each strain. The unpaired *t* test was performed in this figure. *P < 0.05, **P < 0.01, ***P < 0.001. All error bars indicate the mean ± SEM.

**Figure EV6.**
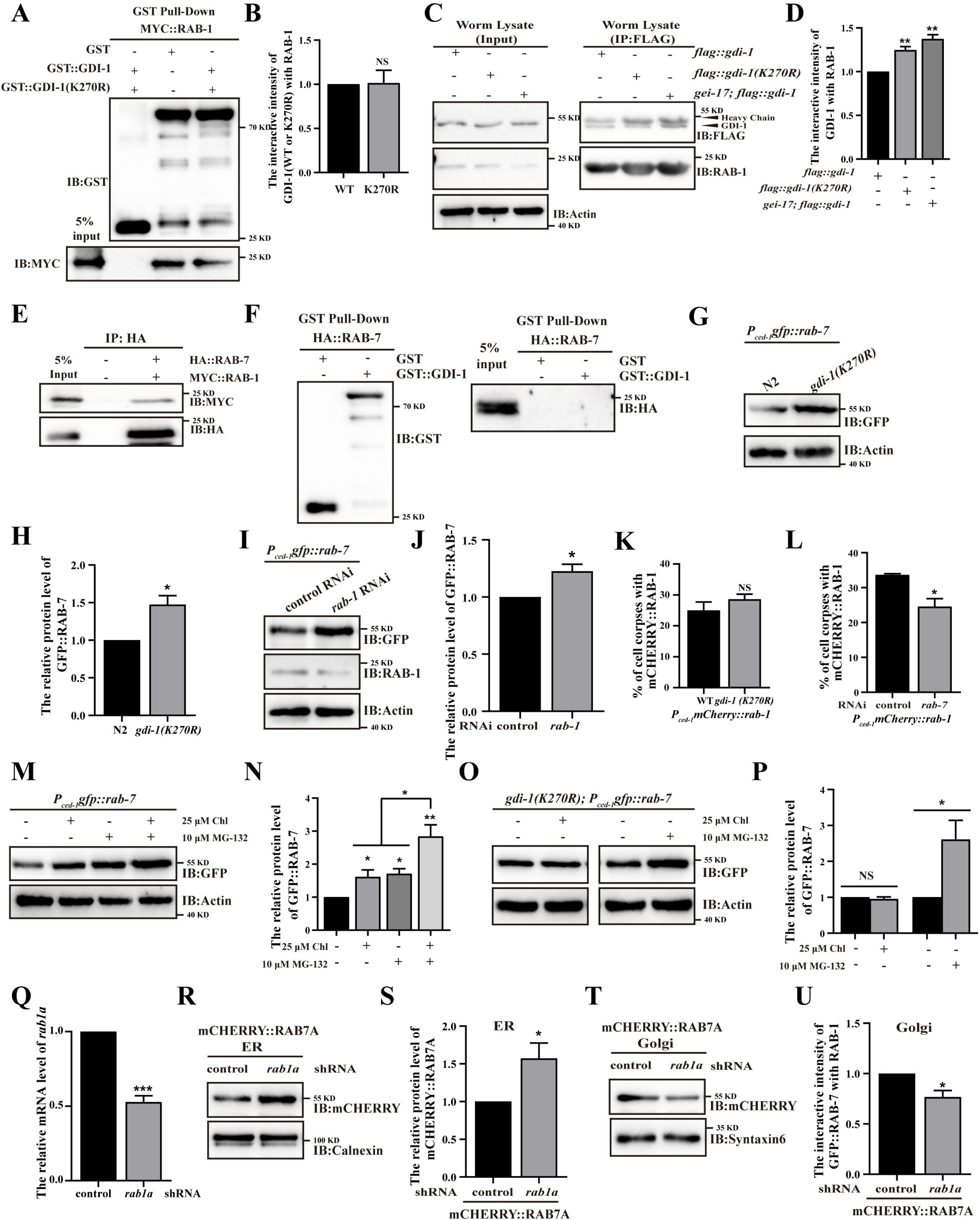
SUMOylation of GDI-1 at K270 regulates RAB-1-mediated vesicular transport of RAB-7 from ER to Golgi. (A and C) The interaction intensity between RAB-1 and GDI-1 (WT/K270R) was examined by the GST pulldown assay (A) and FLAG IP in *flag::gdi-1*, *flag::gdi-1(K270R)*, and *gei-17; flag::gdi-1* worms (C). (B and D) The graphs show the protein levels of RAB-1 and GDI-1. (E) The interaction between RAB-7 and RAB-1 was examined by CO-IP in HEK293T cells. (F) The interaction between RAB-7 and GDI-1 was examined by the GST pulldown assay. (G and H) The graphs shows the percentage referred to the ratio of germ cell corpses labeled by mCHERRY::RAB-1 in *P_ced-1_mcherry::rab-1* and *gdi-1(K270R); P_ced-1_mcherry::rab-1* (G) and *P_ced-1_mcherry::rab-1* control/*rab-7* RNAi (H) worms. (I and K) Exogenous RAB-7 protein levels were examined by immunoblot analysis in *P_ced-1_gfp::rab-7* and *gdi-1(K270R); P_ced-1_gfp::rab-7* (I) and *P_ced-1_gfp::rab-7* control/*rab-1* RNAi (K) worms. (J and L) The graphs show quantification of the protein level of RAB-7 in *P_ced-1_gfp::rab-7* and *gdi-1(K270R); P_ced-1_gfp::rab-7* (J) and *P_ced-1_gfp::rab-7* control/*rab-1* RNAi (L) using ImageJ. (M and O) Exogenous RAB-7 protein levels were examined by immunoblot analysis in *P_ced-1_gfp::rab-7* (M) and *gdi-1(K270R); P_ced-1_gfp::rab-7* (O) treated with 25µM Chloroeuine (lysosome inhibitor) or 10µM MG-132 (proteasome inhibitor). (N and P) The graphs show quantification of the protein level of RAB-7 in *P_ced-1_gfp::rab-7* (N) and *gdi-1(K270R); P_ced-1_gfp::rab-7* (P) treated with 25µM Chloroeuine or 10µM MG-132 using ImageJ software. (Q) *rab1a* mRNA in 293T cells treated with control or *rab1a* shRNA were determined by qRT-PCR. (R and T) Exogenous RAB7A protein levels wereexamined by immunoblot analysis in 293T cells treated with control or *rab1a* shRNA in the ER (R) or Golgi (T). Calnexin is the ER marker; Syntaxin6 is the Golgi marker. (S and U) The graphs show quantification of the protein level of RAB7A in ER (S) and Golgi (U) using ImageJ. The unpaired *t* test was performed in this figure. *P < 0.05, **P < 0.01, NS, no significance. All error bars indicate the mean ± SEM.

**Figure EV7.**
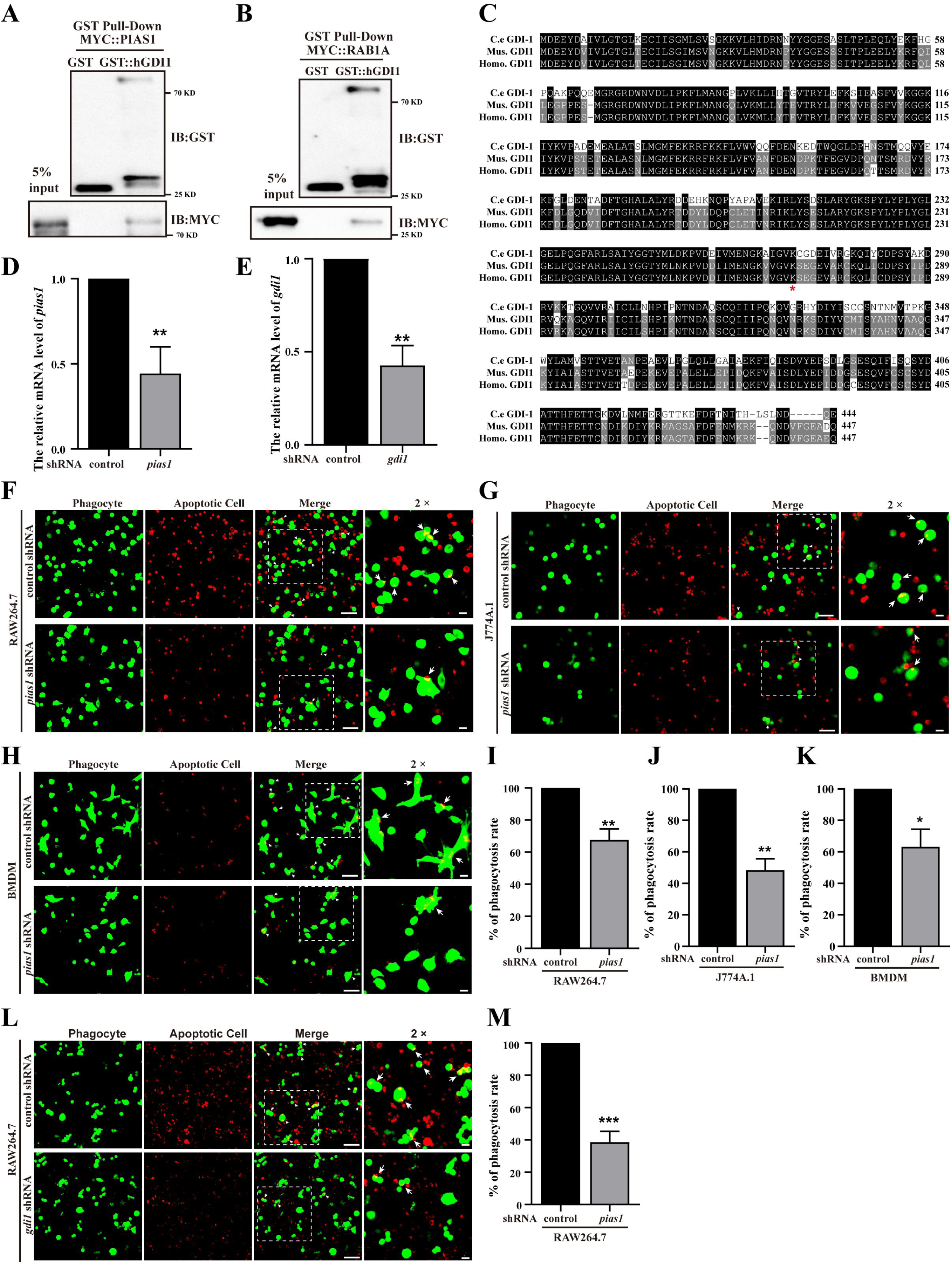
GDI1 SUMOylation regulates efferocytosis in mammals. (A and B) The interaction between hGDI1 and PIAS1 (A) and hGDI1 and RAB1A were examined by GST pulldown assays. (C) Sequence alignment of *C. elegans* (C.e), mice (Mus.), and human (Homo.) GDI-1. Identical residues are shaded in black, and similar ones in gray. The asterisk indicates 270 lysine of C.e GDI-1. (D and E) *pias1* (D) and *gdi1* (E) mRNA in RAW264.7 cells treated with control or *pias1* shRNA (D) and control or *gdi1* shRNA (E) were determined by qRT-PCR. (F-H) Representative images of control and *pias1* shRNA-treated RAW264.7 (F), J774A.1 (G), and BMDM (H) phagocytes (green) engulfing apoptotic Jurkat cells (red). Bars, 100 µm. Arrows indicate the phagocytosis of apoptotic cells. The boxed region is magnified (2×) in the insets (bars, 25 µm). (I-K) Quantification of the percentage of control or *pias1* shRNA treated RAW264.7 (I), J774A.1 (J), and BMDM (K) phagocytes performing apoptotic Jurkat cells engulfment from three independent assays. (L) Representative images of control and *gdi1* shRNA-treated RAW264.7 phagocytes (green) engulfing apoptotic Jurkat cells (red). Bars, 100 µm. Arrows indicate the phagocytosis of apoptotic cells. The boxed region is magnified (2×) in the insets (bars, 25 µm). (M) Phagocytosis rates of control and *gdi1* shRNA-treated RAW264.7 phagocytes performing apoptotic Jurkat cells engulfment from three independent assays. The unpaired *t* test was performed in this figure. *P < 0.05, **P < 0.01, ***P < 0.001. All error bars indicate the mean ± SEM.

